# Inorganic phosphate in Arp2/3 complex acts as a rapid switch for the stability of actin filament branches

**DOI:** 10.1101/2025.07.18.665541

**Authors:** Jiu Xiao, Rebecca Pagès, Guillaume Romet-Lemonne, Antoine Jégou

## Abstract

The rate at which actin filaments turn over modulates actin network architecture and mechanosensitivity. Actin filaments often appear as branches, nucleated by the Arp2/3 complex. Within the Arp2/3 complex, Arp2 and Arp3 are ATPases, which, similar to actin, may adopt slightly different conformations depending on their bound nucleotide. We investigate the impact of the nucleotide state of Arp2/3 complexes on branch stability, by mechanically pulling on them. Remarkably, we find that branches with mammalian ADP-Pi and ADP-Arp2/3 complexes detach with the same exponential increase as a function of force, but ADP-Pi-Arp2/3 complex branch junctions are 20 times more stable at all forces. We observe that inorganic phosphate (Pi) is in rapid equilibrium with ADP-Arp2/3 at the branch junction, and is released at a rate greater than 1 s^-1^. Upon branch dissociation, the surviving Arp2/3 complex, remaining attached to the mother filament, is a thousand times more stable in the ADP-Pi than in the ADP state. Surprisingly, branch regrowth from surviving ADP-Pi-Arp2/3 complex does not require reloading ATP, indicating that the Arp2/3 complex remained in an active state upon debranching. We reveal that GMF destabilizes ADP-Arp2/3 complex branch junctions by accelerating first the dissociation of the daughter filament, and then the dissociation of the surviving Arp2/3 complex from the mother filament. We also report that cortactin stabilizes branches in a force dependent manner, and enhances branch renucleation. GMF and cortactin do not bind to Arp2/3 in the ADP-Pi state. Overall, Our results give a new importance to cytoplasmic Pi, as a possible regulator of branched actin network stability.

## Introduction

Cells assemble a diverse array of actin filament networks to execute spatially and timely controlled tasks throughout the life cycle, including cell motility, cytokinesis, or the transport of vesicles within the cell. The assembly and disassembly of actin filaments are tightly regulated by tens of regulatory proteins. The Arp2/3 complex, composed of seven subunits, including two actin-related proteins, Arp2 and Arp3, stands out as an essential nucleator of actin filaments, uniquely capable of generating branched structures (reviewed in (*1*, *2*)).

The nucleation of actin filament branches by Arp2/3 is a multi-stage process. First, two WASP-family nucleation promoting factors (NPFs) interact with Arp2/3 through their Central and Acidic domains, bind two actin monomers via their WH2 domains, and bring them into contact with Arp2 and Arp3 (*1*, *3–5*). This first step induces allosteric changes leading to the positioning of Arp2 and Arp3 in a side-by-side conformation that resembles the short-pitch actin filament conformation. For its final full activation, this NPF-Arp2/3-actin complex must then bind to the side of a pre-existing actin filament (hereafter referred to as the ‘mother filament’) and the two NPFs must detach from the Arp2/3-actin complex. This final step allows the growth of a filament branch (hereafter referred to as the ‘daughter filament’) by the addition of actin subunits to the newly created barbed end of Arp2/3-actin. This activation of the Arp2/3 complex is accompanied by a flattening of Arp2 and Arp3 subunits (*1*, *5*), analogous to the transition of actin from its globular (G-actin) to filamentous (F-actin) form upon incorporation into a filament (*6*).

To nucleate a branch and be activated by NPFs, Arp2 and Arp3 need to have ATP bound to their nucleotide pocket (*7*). Earlier in vitro biochemical studies on bovine or Acanthamoeba Arp2/3 complex showed that ATP hydrolysis within Arp2 alone accompanies branch formation (*7–9*). Though, in yeast or drosophila, Arp2/3 complexes with either an Arp2 or Arp3 mutant defective for ATP hydrolysis are still competent for branch formation, but not with both (*10*, *11*). This points to an intricate relation between the nucleotide state of the Arp2/3 complex and its activation. In addition, these ATP defective mutations also decrease the stability of branches they form, further highlighting the importance of the nucleotide inside the Arp2/3 complex at branch junctions. The structure of the Arp2/3 complex at a branch junction has been extensively characterized using cryo-electron microscopy (cryo-EM) (*12–15*). For the bovine Arp2/3 complex, cryo-EM maps of mature branches indicate that both Arp2 and Arp3 have an ADP nucleotide bound in their nucleotide-binding pocket (*12*). The Arp2/3 complex thus transitions from the ADP-Pi state towards the ADP state by releasing Pi. ADP-Arp2/3 complex branch junctions have been reported to be less stable than the ADP-Pi-Arp2/3 complex ones (*8*, *9*, *16*), although this difference was only visible in the presence of a pulling force (*16*). Moreover, how fast Pi is released from Arp2/3 complex has remained challenging, with debated rates, depending on the Arp2/3 complex species and the experimental methods employed, varying from seconds for Acanthamoeba Arp2/3 complexes (*7*), to minutes for bovine (*8*) or S. Pombe yeast (*16*) Arp2/3 complexes. The nucleotide state for actin plays a central role in tuning the kinetics rate of filament assembly/disassembly, as well as how regulatory proteins interact with actin subunits of a filament (*17*). Thus, one may expect that the nucleotide state of the Arp2/3 complex at the branch junction should similarly influence the mechanical stability of branches, and how proteins such as Glia Maturation Factor (GMF), cortactin, or coronin (reviewed in (*2*, *18*)) binds to Arp2 or Arp3 subunits at the branch junction, with important consequences on the overall architecture and dynamics of branched actin networks. To investigate this lingering question, one needs to take into account the impact of mechanics on branch stability and the ability of Arp2/3 complexes to renucleate a branch upon branch dissociation, as recently evidenced in vitro by our group (*19*). Though challenging, the combination of these results would provide valuable information to gain insights into branch aging and turnover.

In this study, we investigate the mechanical stability of mammalian Arp2/3 complex branch junctions as a function of their nucleotide state (Fig. 1-4), both in the presence and absence of regulatory proteins, human GMFγ (Fig. 5) or mouse cortactin (Fig. 6). We find that inorganic phosphate (Pi) is in rapid equilibrium with the Arp2/3 complex at the branch junction, and significantly enhances branch stability. Following branch detachment, the dissociation of the surviving Arp2/3 complex in the ADP-Pi state from the mother filament is two orders of magnitude slower than that in the ADP state, further increasing the probability to regrow a branch (Fig. 4). Furthermore, the ADP-Pi state of Arp2/3 complex efficiently prevents GMF-induced debranching as well as cortactin-induced protection. We propose that inorganic phosphate acts as a rapid molecular switch that controls the turnover of branched actin networks within cells (Fig. 7).

**Figure 1.**
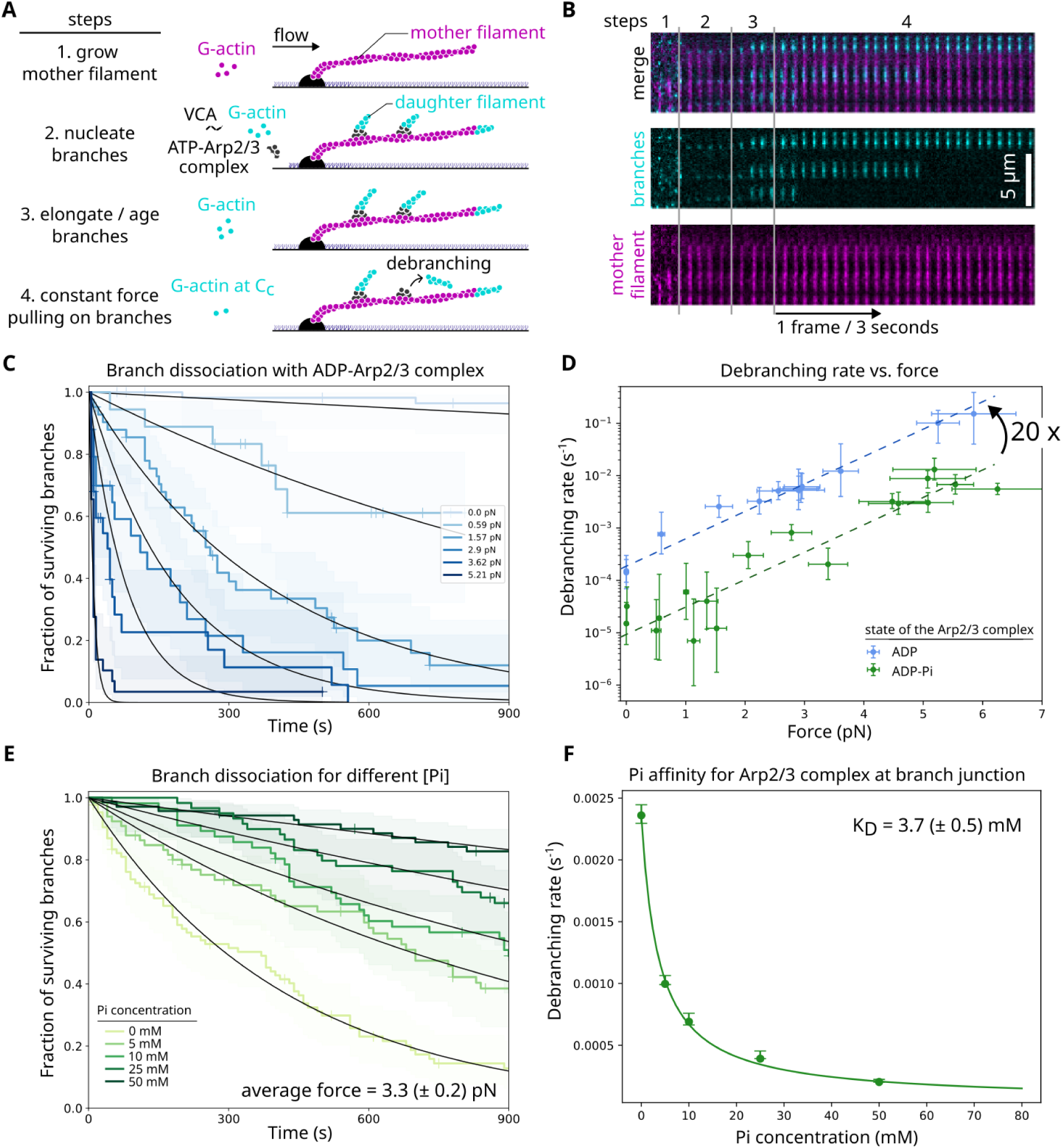
Stability of Arp2/3-mediated actin filament branches probed by constant force pulling. A. Schematic of the debranching experiment using microfluidics. Mother filaments are elongated from surface-anchored spectrin-actin seeds by flowing in 0.7 µM Alexa Fluor 488 (10%)–G-ATP-actin (magenta) for 5 minutes. Branches are nucleated by flowing in 0.4 µM Alexa Fluor 568 (10%)–G-ATP-actin (cyan), with VCA domains from N-WASP, and bovine Arp2/3 complex. They are elongated for 30 minutes by flowing in 0.2 µM Alexa Fluor 568 (10%)–G-ATP-actin alone. The dissociation of branches under constant pulling force are then monitored over time by flowing in 0.1 µM Alexa Fluor 568 (10%)–G-ATP-actin at different flow rates. The flowing solution applies a pulling force to the mother filaments and the branches. B. Time lapse showing the different steps, described in panel A, for branch formation and dissociation, with the actin mother filament is in magenta, and actin branches in cyan. The scale bar is 5 µm. C. Surviving fractions of branches monitored over time using different flow rates to apply different constant forces, while flowing in 0.15 µM Alexa Fluor 568 (10%)–G-ATP-actin. Black curves are single exponential fits. (number of analyzed branches = 60 (0 pN), 18 (0.59 pN), 42 (1.57 pN), 25 (2.9 pN), 25 (3.62 pN), and 29 (5.21 pN)). Shaded areas are 95% confidence intervals. Vertical ticks on the surviving curves are times where censoring events occurred. D. Debranching rates as a function of constant pulling forces, for ADP-Arp2/3 complex branch junctions (blue), or for ADP-Pi-Arp2/3 complex branch junctions by exposing them to a 50 mM Pi buffer solution (green). Error bars are standard deviations for the force, and confidence intervals on the single exponential fits, obtained by fitting the upper and lower bounds of 95% confidence intervals of the survival fractions of debranching experiments. Data points at 0 pN are from experiments performed in ‘open’ chambers, without microfluidics flow. Data obtained from experiments with on average 40 and 100 analyzed branches for ADP- and ADP-Pi-Arp2/3 complex, respectively. Dashed lines are single exponential fits. E. Surviving fractions of branches monitored over time by applying an average force of 3.3 (± 0.2) pN, and exposing them to buffers of different Pi concentrations, containing 0.15 µM Alexa Fluor 568 (10%)–G-ATP-actin. Black curves are single exponential fits. (number of analyzed branches = 86 (0 mM Pi), 66 (5 mM Pi), 56 (10 mM Pi), 70 (25 mM Pi), and 60 (50 mM Pi)). Shaded areas are 95% confidence intervals. Vertical ticks on the surviving curves are times where censoring events occurred. F. Debranching rates as a function of Pi concentration obtained from fit from curves shown in (E). Error bars are confidence intervals on the single exponential fits, obtained by fitting the upper and lower bounds of 95% confidence intervals in panel E.

## Results

### Branch dissociation rate depends on Arp2/3 nucleotide state

To investigate the mechanical stability of Arp2/3-mediated actin filament branches, we employed an in vitro microfluidics-based assay (*19*, *20*). Actin filaments are elongated from spectrin-seeds anchored to the glass surface, and branches are initiated and grown by exposing filaments to a solution containing the VCA domains of N-WASP, Arp2/3 complex, and actin (Fig. 1A, B). Branches are then exposed to a flow solution to apply a viscous drag force and pull on them in a controlled manner (*21*). All experiments were carried out using bovine Arp2/3 complex, with rabbit alpha-skeletal muscle actin in buffer at pH 7, at 25°C (see Methods section).

First, we examined the destabilization of ADP-Arp2/3 complex branch junctions by applying constant pulling forces on them. Given the variability in the reported ATP hydrolysis and Pi release rates for Arp2/3 complexes, we chose to probe the detachment of daughter filaments from branches aged for 30 minutes to ensure Arp2/3 complexes were in the ADP state (Fig. 1A). For each constant force experiment, the fraction of intact branches was fitted with a single exponential function to derive the debranching rate at that specific force (Fig. 1C). For all forces, the surviving curves exhibited a clear single exponential behavior, indicating a constant detachment rate over time. Furthermore, the debranching rate increased exponentially with the applied pulling force (Fig. 1D) which indicates a slip bond behaviour of the rupturing Arp2/3 interface leading to debranching. When branches were aged for 4 minutes only, similar debranching rates were obtained (fig. S1), meaning that either all branches are already ADP after 4 minutes, or that ADP- and ADP-Pi-Arp2/3 complex branch junctions detach with the same rate.

We next investigated Arp2/3 complex branch stability in phosphate buffers of increasing phosphate concentration at a given pulling force (3.3 pN). The debranching rate gradually decreased with increasing phosphate concentration (Fig. 1E), with an equilibrium dissociation constant (K_D_) for Pi binding to ADP-Arp2/3 complex at a branch junction of 3.7 (± 0.5) mM (Fig. 1F). In the presence of a saturating amount of phosphate (50 mM), the detachment rate of branches increased exponentially with force, similarly to the ADP-Arp2/3 complex case, but the rate was systematically ∼ 20 fold lower at all forces (Fig. 1D), consistent with previous observations for S.Pombe yeast Arp2/3 complex branch junctions (*16*). Contrary to this latter study, we observed that the ∼ 20 fold difference in debranching rates between ADP- and ADP-Pi-Arp2/3 complex branch junctions is maintained in the absence of pulling force (Fig. 1D), by performing debranching experiments in standard ‘open’ chambers (without microfluidics, fig. S2).

Beryllium fluoride (BeFx) is a structural analog of inorganic phosphate, that have been classically used to probe actin filament assembly mimicking the ADP-Pi- or ATP-actin state (*22–24*), and to investigate the different structural conformations of actin subunits within filaments based on their nucleotide state (*25*). Notably, BeFx remains stably bound within the F-actin nucleotide pocket for hours (*22*). It was previously shown that BeFx stabilizes S. Pombe Arp2/3 complex branch junctions (*16*). Recent cryoEM observations by Oosterhert and colleagues did not report any BeFx bound on the surface of actin filaments (*25*). We thus assume that BeFx would bind only within the nucleotide pocket of Arp2 and Arp3, not on the surface of Arp2/3 complexes. We thus sought to investigate the force response of Arp2/3 complex branch junctions in the ADP-BeFx state, as a tool to get a stable ADP-Pi-like Arp2/3 complex at the branch junction.

Branches were formed with ATP-Arp2/3 complexes and aged in the presence of 2 mM BeFx for 6 minutes prior to force-induced debranching in a regular buffer that does not contain BeFx (Fig. 2A). We observed debranching rates for ADP-BeFx-Arp2/3 complex branch junctions similar to junctions with ADP-Pi-Arp2/3 complexes (Fig. 2B). We also observed that debranching was not affected by the presence or absence of BeFx in solution (fig. S3). As an additional control, branches exposed to a buffer containing 2 mM sodium sulfate behave similarly to branches in the ADP state (fig. S4), indicating that ions with similar stabilizing protein capacity play no role in stabilizing branch junctions. These observations further support our interpretation that the slower debranching observed in phosphate buffer is primarily due to Pi incorporation into Arp2/3 complexes, rather than a general change in the electrostatic environment of the branch junction or the actin filaments.

**Figure 2.**
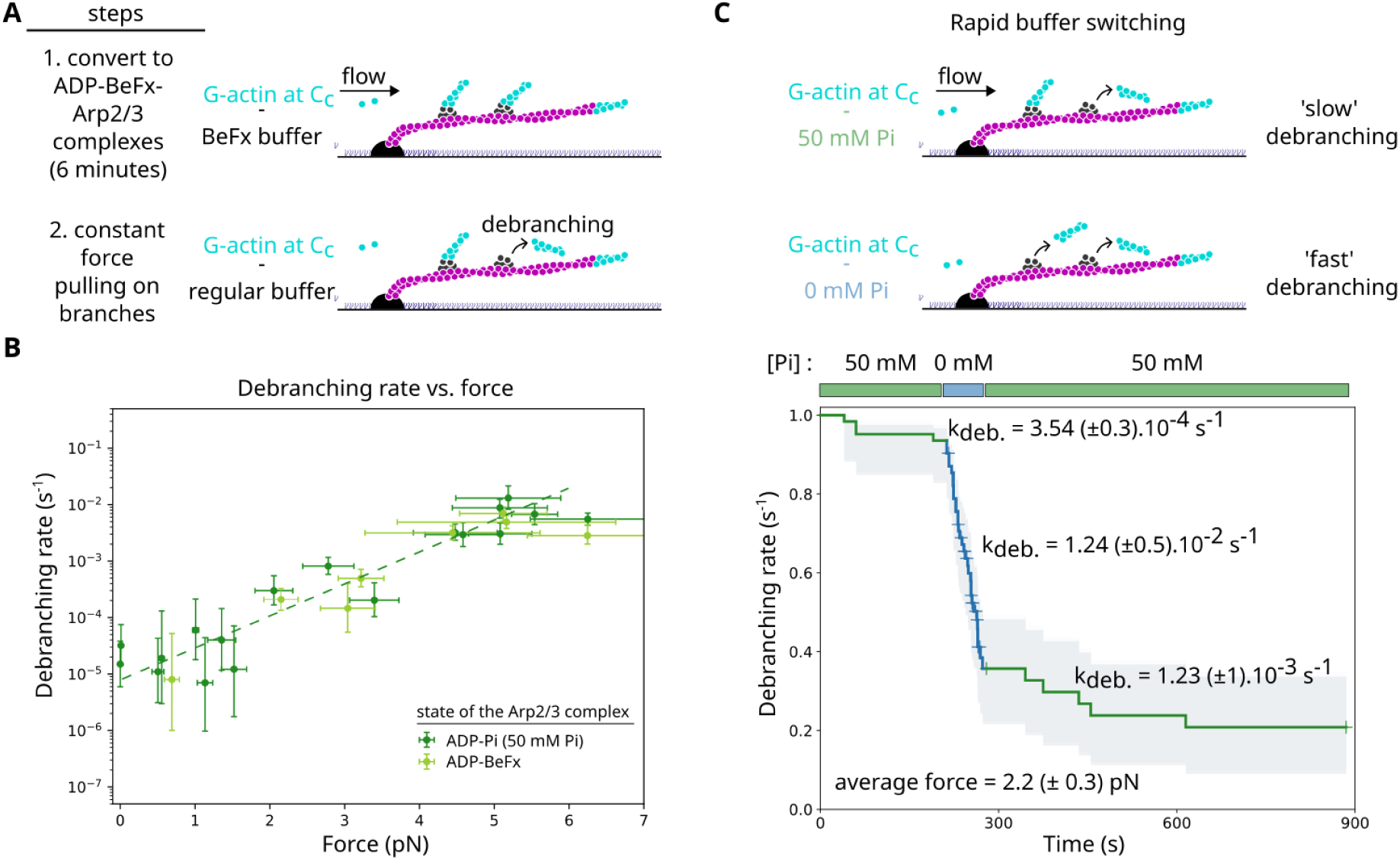
Inorganic phosphate is in rapid equilibrium with the Arp2/3 complex at the branch junction. A. Schematic of the debranching experiment for ADP-BeFx Arp2/3 complex branch junctions. Branches initiated as detailed in Fig. 1A are exposed to a solution containing 2 mM BeFx for 6 minutes. Debranching events are recorded upon switching to a regular buffer solution containing only 0.1 µM Alexa Fluor 568 (10%)–G-ATP-actin at different flow rates. B. Debranching rates as a function of constant pulling forces, for ADP-Pi-Arp2/3 complex branch junctions by exposing them to a 50 mM Pi buffer solution (green), and for ADP-BeFx-Arp2/3 complex branch junctions (light green). The dashed line is a single exponential fit of the ADP-Pi state data. Error bars are standard deviations for the force, and confidence intervals on the single exponential fits, obtained by fitting the upper and lower bounds of 95% confidence intervals of the survival fractions of debranching experiments. For each force condition, data are from experiments with on average 100 and 90 analyzed branches for ADP-Pi- and ADP-BeFx-Arp2/3 complexes respectively. C. Rapid switching between 50 mM Pi or regular buffer leads to rapid change of the debranching rate. Surviving fraction of branches monitored over time by applying an average force of 2.27 (± 0.3) pN and exposing them to buffers always containing 0.1 µM Alexa Fluor 568 (10%)–G-actin. Debranching rates obtained from exponential fits (not shown) of the different phases of the curves are indicated. Around the time at which buffers were rapidly changed by microfluidics, the acquisition frame rate was 2 frames per second. Vertical ticks on the surviving curves are times where censoring events occurred. n = 62 analyzed branches.

Overall, our data reveal that ADP- and ADP-Pi-Arp2/3 branch junctions have debranching rates with similar exponential responses to force. However, the presence of inorganic phosphate inside Arp2 and Arp3 strongly stabilizes branch junctions, over a broad range of pulling forces, including in the absence of force.

### Inorganic Phosphate Readily Associates and Dissociates from Arp2/3 complexes at Branch Junctions

Having shown that inorganic phosphate stabilizes Arp2/3 complex branch junctions with millimolar affinity, we next investigated the kinetics of Pi release from Arp2/3 complexes. We used our microfluidics system to change the solution surrounding the filament branches in approximately 0.5 second (fig. S5) (*26*), while monitoring branch dissociation at a rate up to 10 frames per second. Following branch formation, we recorded their dissociation in a 50 mM phosphate buffer before rapidly switching to a regular buffer containing no phosphate, and then switching back to the initial 50 mM phosphate buffer after a few tens of seconds. Throughout the process, we applied a constant 2.2 pN pulling force. For each phase of the experiment, branches detached at rates similar to either ADP-Pi- or ADP-Arp2/3 complex branch junctions (Fig. 2C), with a ∼ 20-fold difference between the rates observed in the two phases, in agreement with above results (Fig. 1D). Upon buffer switching, the survival curve of the branches displays a sharp kink, indicative of an abrupt transition from a rapid to a slow debranching rate (and vice-versa). Given our temporal resolution, we conclude that both rates at which Pi incorporates and departs from Arp2/3 complexes are greater than 1 s^-1^. Our measured Pi release rate from Arp2/3 complex at branch junctions is considerably faster than what has been previously observed for bovine Arp2/3 complexes at branch junctions using ATP crosslinking bulk assays (∼ 0.001 s^-1^ (*8*)). This is also orders of magnitude faster than Pi release from mammalian alpha-skeletal and beta-actin subunits within the core of actin filaments (0.007 s^-1^ (*20*, *27–30*)) (see Discussion).

### The Differential Stability of Arp2/3 Complex-Mother and Daughter Filament Interfaces is Force Dependent

We next wondered how the relative stability of the two Arp2/3 complex interfaces at the branch junction could vary with applied pulling force. Depending on which interface ruptures first and causes the branch to dissociate from the mother filament, the Arp2/3 complex can either remain on the mother filament, and potentially regrow a branch, or dissociate from the mother filament.

To gain insights into interface stability we needed to combine the information obtained from debranching experiments with the information from renucleation experiments, as a function of force. As a simple model, each Arp2/3 complex interface, either with the mother filament or the daughter filament (Fig. 3A), ruptures with a rate that varies with the applied force according to the transition state theory for a ‘slip’ bond (Fig. 3B) (*31*), following the Bell-Evans relation, k_off_(F) = k_off_(F=0) ⋅ exp(F⋅Δx/k_B_T), where Δx is the distance between the bound and transition states along the reaction coordinate, and k_B_⋅T is the thermal energy (4.1 pN.nm at 25°C). The debranching rate is k_deb_(F) = k_off_^daughter^(F) + k_off_^mother^(F). The probability to renucleate a branch is given by the probability that the Arp2/3 complex-daughter interface ruptures first, which can be written as k_off_^daughter^(F) / (k_off_ ^daughter^(F) + k_off_^mother^(F)), under saturating actin and ATP concentrations (as previously established in (*19*)).

**Figure 3.**
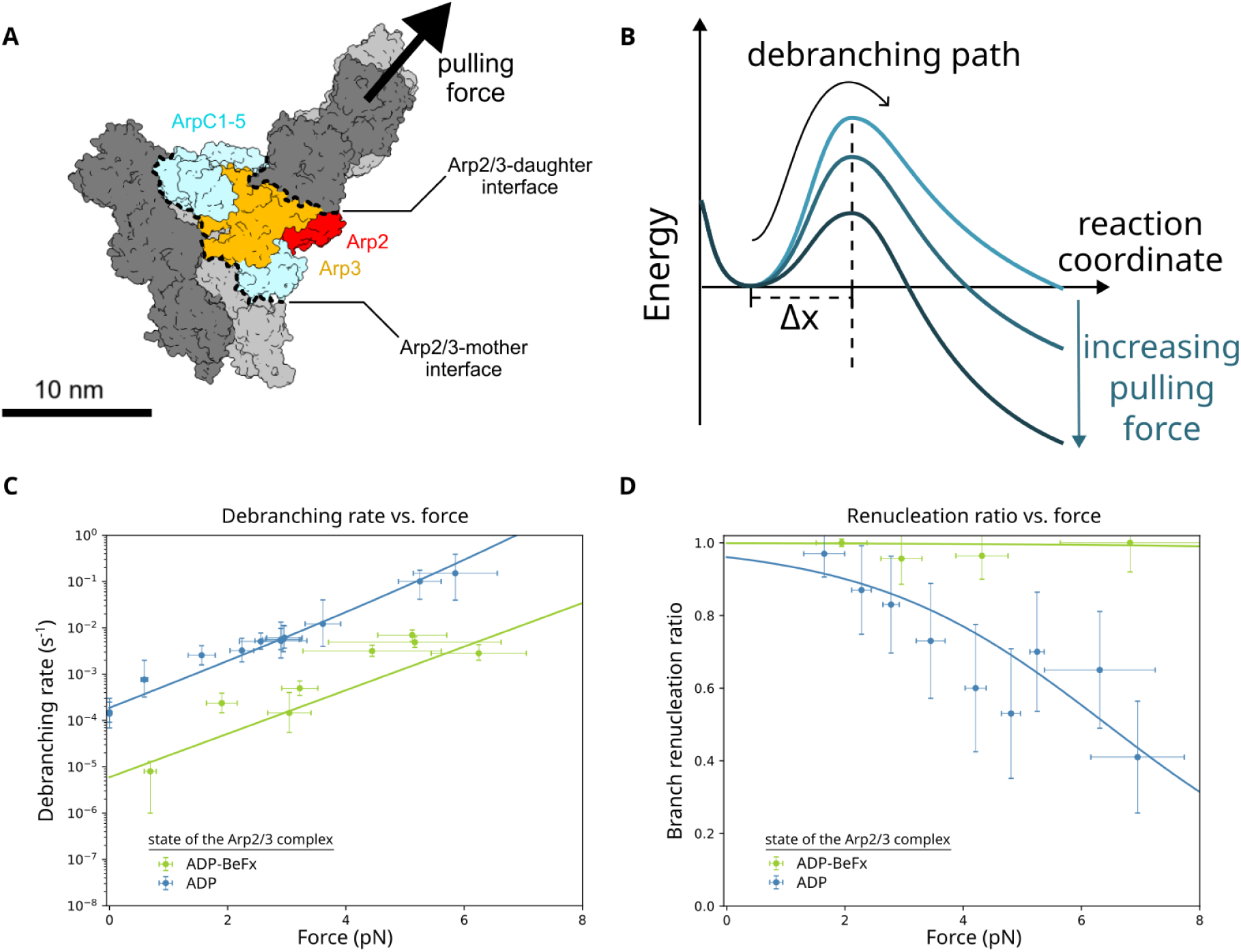
Stability of the Arp2/3 complex interfaces is nucleotide dependent. A. Schematic of the Arp2/3 complex interfaces, adapted from PDB 7TPT (*12*). B. Schematic description of the energy landscape of the interface of the Arp2/3 complex with either the mother or the daughter filament, for which the energy barrier is lowered as a separating pulling force is applied. C. Debranching rates for different pulling forces for ADP- (blue) or ADP-BeFx- (green) Arp2/3 complex branch junctions, in the presence of 0.5 µM actin. Data obtained from experiments with on average 40 and 90 analyzed branches for ADP- and ADP-BeFx-Arp2/3 complexes, respectively. In panel C and D, lines are best fits of the simultaneous error minimization of the probability of renucleating a branch and of the debranching rate, for each Arp2/3 complex state (see Main text). D. Branch renucleation ratio for different pulling forces for ADP- (blue) or ADP-BeFx- (green) Arp2/3 complex branch junctions, in the presence of 1.5 µM Alexa Fluor 568 (10%)–G-actin and 200 µM ATP. Each point is from a single experiment with at least 30 analyzed branches.

First, we conducted branch renucleation experiments for ADP-Arp2/3 complexes at different pulling forces, in the presence of 1.5 µM actin and 200 µM ATP (Fig. 3D). The simultaneous fit of the debranching rate and the renucleation ratio experimental data (Fig. 3C,D, Table 1) indicated that, at zero force, the ADP-Arp2/3 complex-mother filament interface is ∼24 times more stable than the ADP-Arp2/3 complex-daughter interface. This difference in interface stability diminishes with increasing force. Above 6.4 pN, the Arp2/3 complex-mother interface becomes less stable than the interface with the daughter filament, and the majority of Arp2/3 complexes detach from the mother filament while remaining bound to the branch at the debranching time.

**Table 1.** Parameters of the stability of the Arp2/3 complex interfaces. Obtained from the simultaneous fit of the debranching rate and the branch renucleation ratio as a function of force.

| | Arp2/3 complex-daughter filament interface | | Arp2/3 complex-mother filament interface | | $\frac{k_{off, F=0}^{daughter}}{k_{off, F=0}^{mother}}$ |
| --- | --- | --- | --- | --- | --- |
| branch junction | $k_{off, F=0}^{daughter}$ (s<sup>-1</sup>) | $\Delta x^{daughter}$ (nm) | $k_{off, F=0}^{mother}$ (s<sup>-1</sup>) | $\Delta x^{mother}$ (nm) | |
| **ADP-Arp2/3 complex** | 1.8 10<sup>-4</sup> | 4.65 | 7.4 10<sup>-6</sup> | 6.69 | 24.4 |
| **ADP-BeFx-Arp2/3 complex** | 5.2 10<sup>-6</sup> | 4.51 | 5.8 10<sup>-9</sup> | 5.64 | 907 |
| **cortactin-bound ADP-Arp2/3 complex** | 1.9 10<sup>-4</sup> | 2.82 | 2.2 10<sup>-7</sup> | 5.80 | 850 |

We performed similar experiments with ADP-BeFx-Arp2/3 complex branch junctions (Fig. 3C,D). Strikingly, we observed that the renucleation ratio for ADP-BeFx-Arp2/3 complexes remains high (> 95%) for a range of forces much larger than for ADP-Arp2/3 complex branch junctions. According to the fit, at zero force, the ADP-BeFx-Arp2/3 complex-mother interface is > 100 times more stable than the daughter interface, and becomes less stable than the daughter interface only above 20 pN, a transition force much higher than for Arp2/3 complexes in the ADP state (fig. S6, Table 1). This indicates that the presence of BeFx strengthens both interfaces, rendering the interface with the mother extremely stable.

The renucleation ratio of ADP-BeFx-Arp2/3 complex branch junctions remained high for several generations of renucleated branches, even at low actin concentration (fig. S7), indicating that the Arp2/3 complex remained in the ADP-BeFx state for several rounds of renucleation. This implies that branch renucleation does not require reloading ATP when the Arp2/3 complex is in the ADP-BeFx state. Thus surviving ADP-BeFx-Arp2/3 complexes remain ‘active’, Arp2 and Arp3 stay in a barbed end-like conformation, ready to bind actin monomers to grow a new branch.

This striking observation of the stability of the ADP-BeFx-Arp2/3 complex-mother interface (Fig. 3) prompted us to determine the unbinding rate of surviving ADP-BeFx-Arp2/3 complexes from mother filaments. To investigate this, debranching was performed by pulling on branches in the absence of actin in solution with a moderate force (∼ 5 pN) for a maximum of 13 minutes, before switching to a solution containing 0.4 µM actin in order to reveal the presence of remaining surviving ADP-BeFx-Arp2/3 complexes by renucleating branches from them (Fig. 4A). Surviving ADP-BeFx-Arp2/3 complexes exhibited remarkable stability (Fig. 4B), detaching from mother filaments at a rate of 6.86 10^-4^ s^-1^, almost a 1000 times slower than for ADP- or ATP-Arp2/3 complexes (∼ 0.6 s^-1^, (*19*), see Discussion).

**Figure 4.**
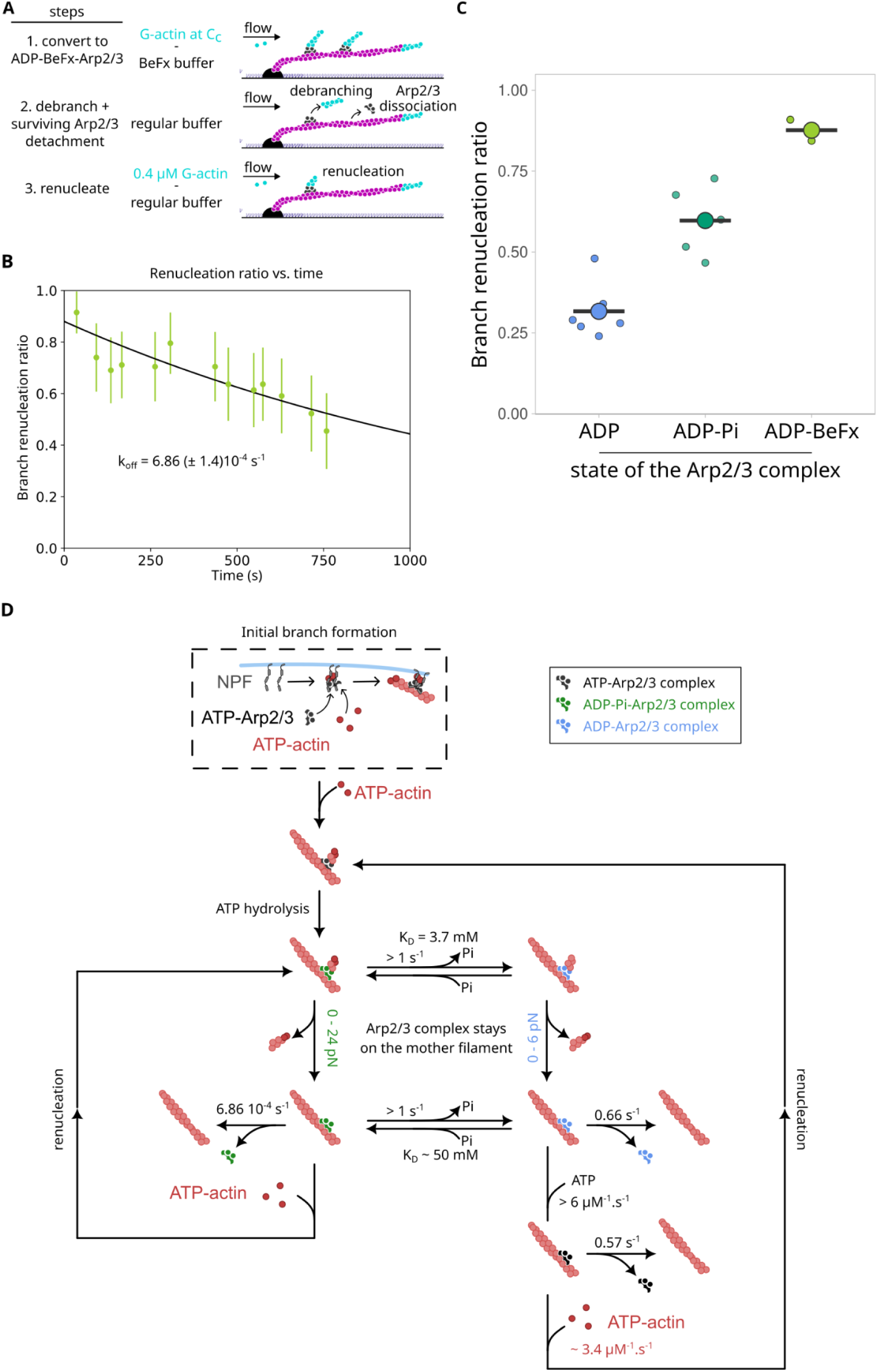
Branch renucleation ratio as a function of Arp2/3 complex nucleotide state. A. Schematics of the renucleation experiment for ADP-BeFx-Arp2/3 complex branch junctions. Branches are nucleated and converted to BeFx for 6 minutes as in Fig. 2A, then exposed to buffer only for 3 (n = 184 branches), 10 (n = 132 branches), or 13 minutes (n = 264 branches), before being exposed again to a F-buffer solution containing 0.4 µM Alexa Fluor 568 (10%)–G-ATP-actin (cyan) to allow branch to regrow. B. Branch renucleation ratio for ADP-BeFx-Arp2/3 complex branch junctions, probed as shown in A, pulling with an average force of 5.2 pN. Each datapoint represents over 40 branches. Error bars are standard deviations. Exponential fit of the data gives a rate of detachment of the surviving ADP-BeFx-Arp2/3 complex from the mother filament of 6.86 (± 1.4) 10^-4^ s^-1^. C. Branch renucleation ratio from ADP-(blue), ADP-Pi-(green) or ADP-BeFx-(light green) Arp2/3 complex branch junctions, in the presence of 0.5 µM Alexa Fluor 568 (10%)–G-actin, pulling with an average pulling force of 4.9 pN. Each individual (small) data point is from a single experiment with at least 30 analyzed branches. Large data points represent the average value for each condition. D. Reaction scheme of the regulation of actin filament branches as a function of force and phosphate. After initial branch initiation by the coordinated action of membrane-bound NPF, actin and inactive ATP-Arp2/3 complex (dashed box), ATP-Arp2/3 quickly becomes ADP-Pi-Arp2/3. Depending on Pi concentration, branches are in rapid equilibrium between ADP- and ADP-Pi states. ADP-Pi-Arp2/3 complex branch junctions are more resistant to pulling force. In both ADP and ADP-Pi cases, at low force regime, the Arp2/3 complex-daughter interface ruptures before the Arp2/3 complex-mother interface. The surviving ADP-Pi-Arp2/3 complex detaches very slowly from the mother filament. Pi affinity for surviving ADP-Arp2/3 complexes is ∼ 10 fold lower than for ADP-Arp2/3 complexes at branch junctions. Renucleation in ADP-Pi state does not require ATP reloading. Rates for the ADP state as from (*19*).

In 50 mM phosphate buffer, where Arp2/3 complexes at branch junctions are predominantly (>95%) in the ADP-Pi state, and in the presence of 0.5 µM actin and 200 µM ATP, the renucleation fraction was intermediate between the ones of ADP-Arp2/3 branches and ADP-BeFx-Arp2/3 branches obtained with the same actin and ATP concentrations (Fig. 4C). This indicates that, once the daughter filament detaches from the Arp2/3 complex, the affinity of the Arp2/3 complex for Pi drops dramatically, and around 55% will become ADP-Arp2/3 complexes by releasing Pi. A surviving ADP-Pi-Arp2/3 complex can therefore dissociate from the mother filament (rare cases), lose its Pi and transition towards the ADP state despite the presence of 50 mM phosphate in solution (∼55% of the cases), or renucleate a branch while staying in the ADP-Pi state (∼45% of the cases). Considering the outcome of the competition between these three alternatives routes, and assuming a binding rate of actin to the surviving ADP-Pi-Arp2/3 complex similar to what we determined previously for ATP-Arp2/3 (*19*), i.e. k_on_ = 3.4 µM^-1^.s^-1^ (see also fig. S6), the Pi release rate from surviving ADP-Pi-Arp2/3 complexes would be of the order of k_r,Pi_ ∼ 1.35 s^-1^. All the reactions are summarized in Fig. 4D.

### GMF destabilizes the ADP-Arp2/3-daughter interface

GMF (Glia Maturation Factor) is a member of the ADF-homology protein family (reviewed in (*32*)). Mammals express two isoforms, GMFβ and GMFγ, sharing 82% sequence identity and strong structural similarities, in a tissue dependent manner. GMF plays an important role in regulating lamellipodia dynamics and controlling cell motility (*33*). Although structurally very similar to other members of the family such as cofilin, GMF does not bind to actin but only interacts with the Arp2/3 complex. When interacting with the Arp2/3 complex in the inactive splayed conformation, GMF binds to Arp2 and ArpC1 (*34*). Moreover, GMF has been reported to preferentially bind soluble Arp2/3 complexes in the ADP state (*35*), reminiscent of the nucleotide dependent affinity of cofilin for actin (*36*). GMF is able to inhibit Arp2/3 nucleation and to also directly promote debranching (*19*, *37–39*). However, the molecular details of how GMF favors debranching and prevents branch renucleation remain unclear. We thus explored how pulling forces and the nucleotide state of the Arp2/3 complex could impact human GMFγ (hereafter referred to as ‘GMF’) activity.

We first exposed ADP-Arp2/3 complex branch junctions to varying concentrations of GMF. First, we observed a mild ∼ 6-fold increase in the debranching rate, with a GMF affinity for ADP-Arp2/3 complex at the branch junction of 30 (± 8) nM (Fig. 5A), consistent with previous studies using *Drosophila* GMF (*19*) or yeast S. Pombe GMF (*16*), but with different potency. Second, the debranching rate of ADP-Arp2/3 complex branch junctions exposed to saturating amounts of GMF (> 500 nM) increases exponentially with force in a similar manner than without GMF (Fig. 5B). Exposing ADP-BeFx-Arp2/3 complex branch junctions to a saturating amount of GMF did not result in an appreciable increase in the debranching rate (Fig. 5B).

**Figure 5.**
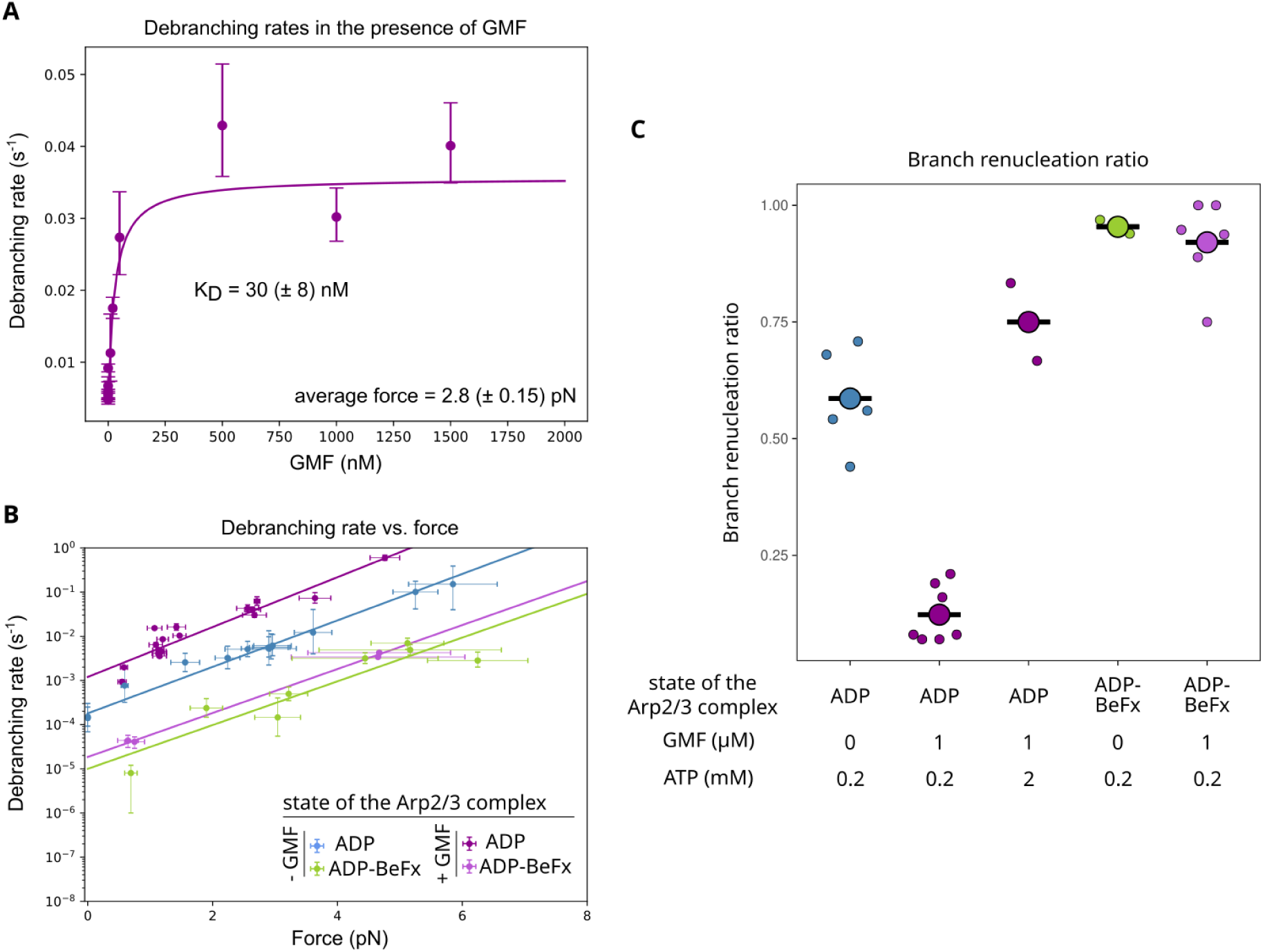
GMF accelerates debranching in a nucleotide dependent manner. A. Debranching rate of ADP-Arp2/3 complex branch junctions exposed to various concentrations of GMF, from experiments performed a constant pulling force (average 2.8 pN), in the presence of 0.15 µM actin. Each data point from a single experiment with at least 30 analyzed branches. Error bars are standard deviations. The binding affinity of GMF for ADP-Arp2/3 complex at the branch junctions derived from the saturation curve fitting is 30 (± 8) nM. B. Debranching rate of ADP- or ADP-BeFx-Arp2/3 complex branch junctions, in the presence or absence of at least 500 nM GMF in solution, for different pulling forces. Continuous lines are exponential fit from the datapoints for each condition. Error bars are standard deviations for the force, and confidence intervals on the single exponential fits, obtained by fitting the upper and lower bounds of 95% confidence intervals of the survival fractions of debranching experiments. Each data point from a single experiment with at least 19 and 50 analyzed branches for ADP- and ADP-BeFx-Arp2/3 complex branch junctions in the presence of GMF. Data in the absence of cortactin are from figure 3. C. Branch renucleation ratio for ADP- or ADP-BeFx-Arp2/3 complex branch junctions, in the presence or absence of at least 1 µM GMF, 0.5 µM actin, and either 200 µM or 2 mM ATP in solution, with a pulling force applied on branches between 1 and 4 pN. Each individual (small) data point is from a single experiment with at least 30 analyzed branches. Large data points represent the average value for each condition (N > 2).

Similar to our previous study with drosophila GMF (*19*), branch renucleation ratio in the presence of GMF was low compared to renucleation in the absence of GMF (Fig. 5C). Unexpectedly, increasing ATP concentration to 2 mM was sufficient to increase branch renucleation ratio in the presence of GMF to levels similar to what was observed for ADP-Arp2/3 complex branch junctions in the absence of GMF (Fig. 5C). This shows that GMF first accelerates debranching by destabilizing the Arp2/3 complex-daughter interface, and, as a second step, accelerates the departure of the surviving Arp2/3 complex from the mother filament. This accelerated departure of the surviving ADP-Arp2/3 complex can be out-competed by increasing ATP concentration to reload ATP inside Arp2/3 complex faster, which allows branch regrowth. As a confirmation that GMF does not appreciably bind to ADP-BeFx-Arp2/3 complexes, even the surviving ones, the branch renucleation ratio for ADP-BeFx-Arp2/3 complexes branches was unaffected by GMF (Fig. 5C). Thus GMF is unable to destabilize ADP-BeFx-Arp2/3 complexes at branch junctions, and unable to accelerate the dissociation of the surviving ADP-BeFx-Arp2/3 complex from the mother filament after debranching. This strongly suggests that GMF is unable to bind to ADP-BeFx-Arp2/3 complexes, despite its ability to bind both to ADP-Arp2/3 complexes at branch junctions (Fig. 5A, and (*37*)) and to Arp2 of inactive ATP-Arp2/3 complexes in solution (*34*).

### Cortactin stabilizes Arp2/3 complex branch junctions in a force dependent manner

Cortactin is a multi-domain protein that regulates cell migration (see (*40*) for a review), and is known to stimulate Arp2/3 activation, in synergy with NPFs (*41*, *42*), by stabilizing the short-pitch active conformation of the Arp2/3 complex (*43*). Notably, cortactin is able to directly target branch junctions by simultaneously binding to Arp3 via its NtA domain, and to the first actin subunits of the daughter filament through its unstructured 6.5 actin binding repeat domain (*15*). This binding mode has been proposed to stabilize branch junctions by preventing the dissociation of the daughter filament from the Arp2/3 complex (*15*, *37*). The cryo-EM structure from Liu and colleagues shows cortactin bound to an ADP-Arp2/3 complex branch junction (*15*), but how cortactin-bound branches respond to mechanical forces, and whether cortactin is sensitive to the nucleotide state of the Arp2/3 complex at the branch junction remain open questions.

We found that the presence of cortactin reduces the debranching rate for ADP-Arp2/3 complex in a force dependent manner (Fig. 6A). At low force (i.e. below ∼ 1 pN), there was no detectable difference in the debranching rate with or without 20 nM cortactin. At 6 pN, cortactin-bound branches were approximately 20 times more stable than without cortactin, reaching the stability of ADP-Pi-Arp2/3 complex branch junctions. Moreover, the branch renucleation ratio of ADP-Arp2/3 complex branch junctions is increased by the presence of cortactin (Fig. 6B). Unexpectedly, this indicates that the detaching interface is not modified by the presence of cortactin and remains the daughter interface for pulling forces ranging from 0 to 9 pN. According to the model fit (Fig. 6 and Table 1), cortactin stabilizes and changes the overall binding energy of the Arp2/3 complex-daughter interface (see Discussion), and strongly reinforces the stability of the ADP-Arp2/3-mother filament interface without affecting its conformation.

**Figure 6.**
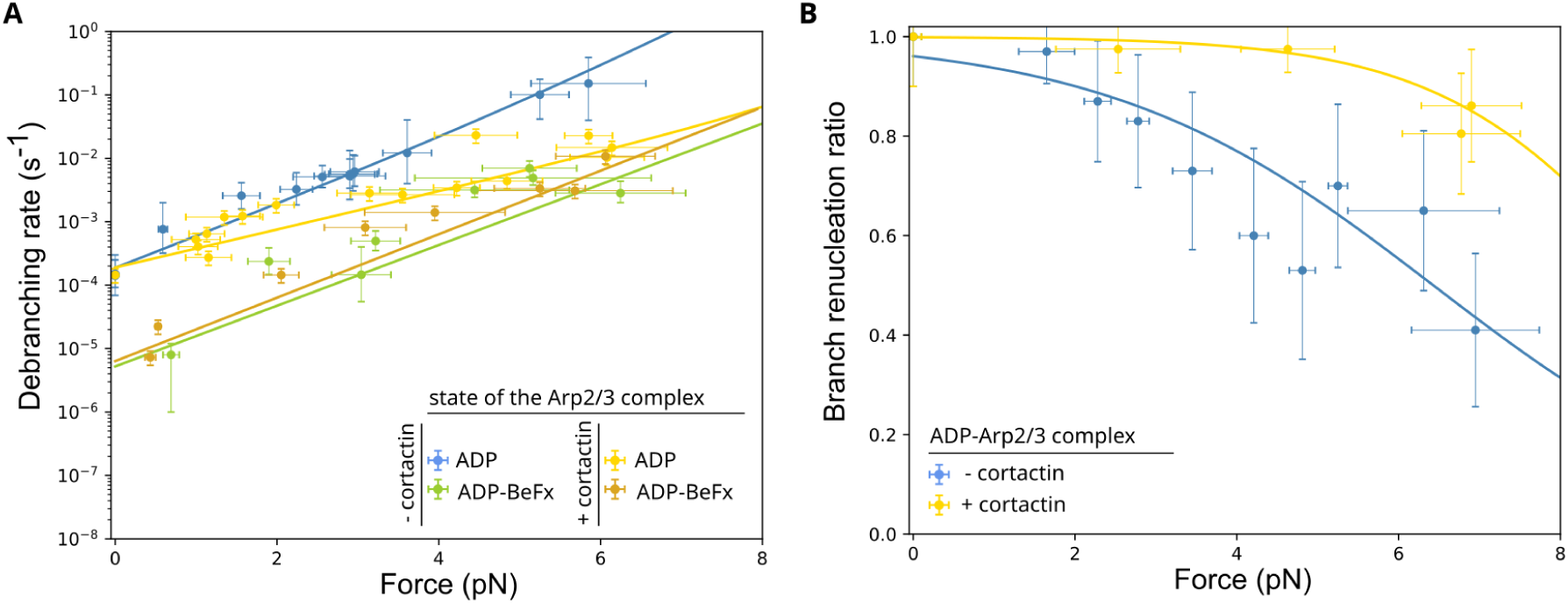
Cortactin stabilizes ADP-Arp2/3 complex branch junctions and favors renucleation. A. Debranching rate for different pulling forces for ADP-(yellow) or ADP-BeFx-(medium gold) Arp2/3 complex branch junctions in the presence of 20 nM cortactin, in the presence of various actin concentrations. Data and fit shown in blue and green are from Fig. 3. Each data point is from a single experiment with at least 60 and 37 analyzed branches for ADP- and ADP-BeFx-Arp2/3 complexes respectively. Error bars are standard deviations. in (A) and (B), lines are best fits of the simultaneous error minimization of the probability of renucleating a branch and of the debranching rate, for each Arp2/3 complex state (see Main text). Data in the absence of cortactin are from figure 3. B. Branch renucleation ratio for different pulling forces for ADP-Arp2/3 complex branch junctions, in the presence (yellow) or absence (blue) of 20 nM cortactin, and 1.5 µM Alexa Fluor 568 (10%)–G-actin and 0.2 mM ATP. Each point is from a single experiment from at least n = 30 branches. Error bars are standard deviations. Data in the absence of cortactin are from figure 3.

Similar to GMF, we did not detect any impact of cortactin on the debranching rate of ADP-BeFx-Arp2/3 complex branch junctions (Fig. 6A), as there was no sign of increased stabilization by cortactin at all forces. We interpreted this as the inability of cortactin to bind to ADP-BeFx-Arp2/3 complex branch junctions.

## Discussion

Here, using an in vitro live microscopy assay, based on microfluidics, with single filament-resolution, we showed that ADP-Pi-Arp2/3 complex branch junctions are around 20 times more stable than ADP-Arp2/3 complex branch junctions, even in the absence of force. This is in contrast with the earlier study on *S. Pombe* Arp2/3 complex, which reported that the difference in branch stability between the two nucleotide states would vanish in the absence of pulling force (*16*). Compared to the rate at which branches dissociate, inorganic phosphate in solution is in rapid equilibrium with the ADP-Arp2/3 complex at the branch junction, with a millimolar binding affinity. We show that, after branch formation, the Arp2/3 complex rapidly (> 1 s^-1^) releases its Pi and becomes ADP. Since Pi can quickly bind back, the branch junctions are in a state, in between ‘young’ and ‘old’, that is tuned by the concentration of inorganic phosphate in solution. As a consequence the Arp2/3 complex branch junction does not have a ‘slow’ actin-like internal clock based on its nucleotide.

Over a physiological range of force (typically 0-5 pN), debranching can be accelerated 200 times with increasing force, the daughter filament detaches from the Arp2/3 complex which remains on the mother filament, and the surviving Arp2/3 complex quickly regrows a branch. This debranching sequence is unaffected by the presence of branch stabilizer cortactin or branch destabilizer GMF, but both proteins are sensitive to the nucleotide state of the Arp2/3 complex at the branch junction. They do not bind to ADP-Pi-Arp2/3 complexes at branch junctions. We have summarized our results in figure 7.

**Figure 7.**
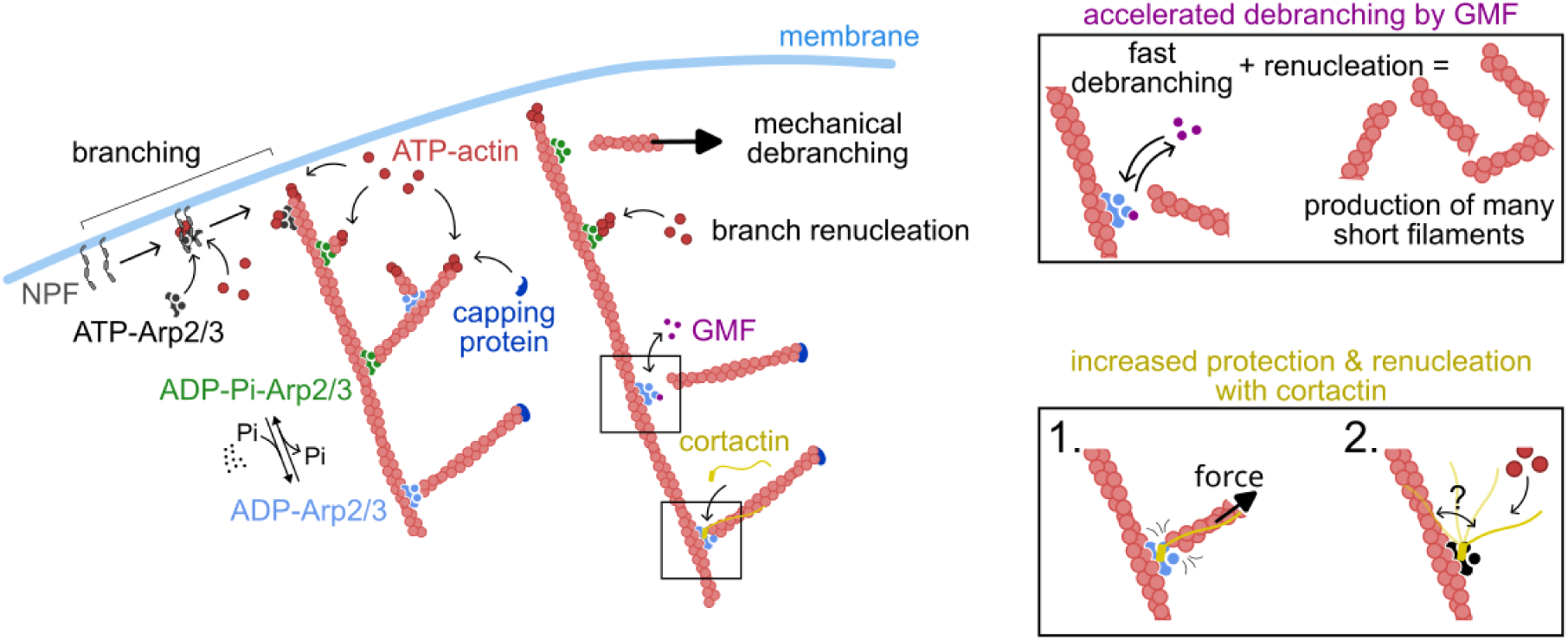
Summarizing sketch of branch regulation by mechanics and regulatory proteins. Actin filaments branches are initiated by membrane bound Nucleation Promoting Factors (NPF) that activate Arp2/3, bind two actin monomers to the pseudo-barbed end formed by Arp2 and Arp3. Upon ATP hydrolysis within the Arp2/3 complex, the cytoplasmic Pi concentration sets the equilibrium between ADP and ADP-Pi states of the Arp2/3 complex at branch junctions. Pulling force accelerates debranching. Branch renucleation is favored by elevated concentration of actin, ATP and Pi in the cytoplasm. GMF and cortactin do not bind to ADP-Pi-Arp2/3 complex. GMF accelerates debranching and dissociation of surviving Arp2/3 (upper inset). Cortactin delays debranching in a force dependent manner, and favors renucleation.

### Molecular Implications

The Arp2/3 complex in the ADP-Pi state is both more stable at the branch junction and as a surviving element of the debranching process, than in the ADP state (Fig. 1, 3, 4). One very striking observation that emerges from this study is that surviving ADP-BeFx-Arp2/3 complexes are much more stable than surviving ADP-Arp2/3 complexes (Fig. 4), or even ATP-Arp2/3 complexes (*19*, *44*). This argues in favor of the ADP-BeFx state of the Arp2/3 complex being different from the ATP state, but mimicking more closely the ADP-Pi state (a question that remains partly unresolved for actin (*22*, *25*)). This would thus indicate that the presence of Pi in Arp2 and Arp3, through allosteric pathways, strongly modulates the stability of the Arp2/3 complex-mother interface. Surprisingly, Chavali and colleagues compared the structures of the branch junction formed by *S. Pombe* Arp2/3 complexes in the ADP and ADP-BeFx states, and reported minor differences ot the Arp2/3 complex interfaces with the mother and daughter filament, with slightly more contacts between Arp2/3 complex and actin subunits in the ADP-Pi state (*45*). These differences puzzlingly may not account fully for the thousand fold difference in stability we observed here. It is thus possible that more than the average position of the residues of the different proteins that compose the branch junction, that seemingly appear very similar in the ADP and ADP-BeFx cases observed in cryoEM, it is in fact the vibration modes of the residues that would tune the binding energy of the interfaces. Pi, and BeFx, would thus play a key role here.

Despite this surprising low dissociation rate, surviving ADP-BeFx-Arp2/3 complexes are still less stable than ADP-BeFx-Arp2/3 complexes at a branch junction under no pulling force (Fig. 1 & 2). The binding of actin to Arp2 and Arp3 thus anchors the Arp2/3 complex to the mother filament, in a tighter or less fluctuating manner.

Inorganic phosphate rapidly shuttles in and out of the Arp2/3 complex (Fig. 2 & 4). According to the conformation of Arp2/3 complex at a branch junction, the backdoor through which Pi is thought to be escaping from the nucleotide pocket in actin (*29*, *30*, *46*) appears wide open in Arp2 and Arp3 (residues N111-R177 in actin, corresponding to N115-R181 in Arp2, and N118-H192 in Arp3, see fig. S8). Moreover, in mammalian Arp2/3, His80 in Arp3 (corresponding to His73 in mammalian actin) is not methylated, and corresponds to residue N77 in Arp3, which is also not modified. It has been established that the absence of methylation on His73 of β-actin accelerates Pi release (*30*). Thus the conformations of Arp2 and Arp3, together with the absence of modification of key residues, can account well for the fast Pi release in Arp2/3 complexes.

Previous studies point towards ATP hydrolysis in Arp2 to have a more central role on branch stability (*10*, *11*), and to take place sooner than in Arp3 (*8*, *9*). Eventually, at least in mammals, both Arp2 and Arp3 have ADP bound in mature branches, as resolved by cryoEM (*12*, *13*, *15*). Our observations do not allow us to identify if Pi binds preferentially to Arp2, Arp3 or both. If the nucleotide state of both Arp2 and Arp3 participate in stabilizing actin branches, Pi release in Arp2 and Arp3 seems to occur both at similar short timescales, as evidenced by our rapid buffer switch experiment (Fig. 2) where the stability of branches are modified in one single fast step upon addition/removal of Pi in solution.

Surprisingly, surviving ADP-BeFx- and ADP-Pi-Arp2/3 complexes do not need to reload ATP to regrow a branch. The presence of Pi maintains some allosteric coordination between residues of the complex, thus Arp2 and Arp3 are able to remain in an ‘active’ short-pitch conformation to allow actin to bind to them and regrow a branch. The fact that the Pi affinity drops by a factor ∼ 10 between the Arp2/3 complex at a branch junction and the surviving ones (Fig. 4) further indicates that Arp2/3 complexes are in different conformations in these two contexts.

At the branch junction, the Arp2/3 complex interfaces are not strongly rearranged by the presence or absence of Pi, as the ADP- and ADP-Pi-Arp2/3 complexes show similar exponentially force-dependent debranching responses (same Δx, Fig. 3 and table 1). They are probably very similar in terms of involved residues making contacts with actin subunits, as confirmed by Chavali and colleagues (*45*). Still, the slight change of conformations upon Pi release is enough for GMF and cortactin to bind (Fig. 5 and 6), probably making key residues more accessible. This is somehow reminiscent of the nucleotide-dependent binding of cofilin to actin, which has been proposed to be due to the higher flexibility of ADP-actin and a more closed D-loop conformation (*25*, *46*).

As revealed in our debranching assay, cortactin binding to ADP-Arp2/3 complex branch junctions strongly modifies the mechanical stability of the rupturing interface upon branch dissociation. Cortactin, being like a leash strongly attached to the daughter filament, adds an additional link between the daughter filament and the Arp2/3 complex (Fig. 7). Upon debranching events, when the Arp2/3 complex-daughter filament interface ruptures, this leash would favor the rebinding of the daughter filament by preventing the daughter to diffuse away quickly (*47*). This would explain the observed increase in branch survival compared to the absence of cortactin. As the pulling force is increased, this rebinding mechanism becomes less efficient. This would reveal that the weakest link is between the NtA domain of cortactin and Arp3. Upon debranching, cortactin would thus detach from the Arp2/3 complex but remain firmly attached to the daughter filament. A new cortactin would then bind to the surviving Arp2/3 complex, strengthening the Arp2/3 complex attachment to the mother filament, leading to an increased renucleation ratio compared to the one of ADP-Arp2/3 complex without cortactin. Cortactin potently protects branches from GMF destabilizing activity (*37*). In a mechanical context, it is hard to anticipate how our proposed ‘leash-protecting’ mechanism of cortactin could prevent GMF from directly binding Arp2.

We now reveal that GMF destabilizes the Arp2/3 complex-daughter interface, leaving the Arp2/3 complex attached to the mother filament (Fig. 5, 7). This contradicts conclusions from recent studies, including ours (*19*, *48*), where it was hypothesized that GMF would induce the departure of the Arp2/3 complex with the branch. Additionally, we now show that GMF is efficient at targeting surviving Arp2/3 complexes, only in the ADP state. To be efficient, GMF-induced dissociation of surviving ADP-Arp2/3 has to happen faster than reloading ATP inside Arp2/3. This might explain why it is extremely difficult to observe the sequential departure of the daughter filament and then of the surviving Arp2/3 complex using single molecule fluorescence approaches (*48*). Furthermore, even in the presence of a saturating amount of GMF in solution, high concentration of ATP allows the exchange of ADP to ATP to out-compete GMF binding to surviving ADP-Arp2/3, and thus lead to branch renucleation.

### Cellular implications

In cells, besides elevated ATP and actin concentrations in the cytoplasm, free inorganic phosphate could play an important role in regulating actin filament branch stability. We showed that Pi affinity for Arp2/3 complexes at branch junctions is around 3.7 mM (Fig. 1), a value which lies within the reported 1-10 mM Pi concentration measured in the cytosol in different mammalian cell types (*49*, *50*) (note that in yeast, Pi concentration is higher, around 10-75 mM (*51*)). Interestingly, even though the release of Pi is fast (∼ 1 second, Fig. 2), it may still be a ‘slow enough’ process for branch junctions to ‘age’, i.e. evolve from ADP-Pi to ADP states, in the case of branched actin networks that turnover very fast. One such case are endocytic patches, made of branches that are nucleated and disassembled within ∼ 1-2 seconds (*52*, *53*).

The ability for the Arp2/3 complex to rapidly switch back towards the ADP-Pi state could provide branches a protective mechanism against GMF debranching activity. The debranching rate can be decreased by 120-fold between the case where debranching is caused by GMF acting on an ADP-Arp2/3 complex branch junction versus when the branch is in the ADP-Pi state in the presence of high Pi concentration (Fig. 4). This holds true over a large range of force (0 to 6 pN).

Unexpectedly, cortactin, which is thought to be a branch stabilizer, also appears unable to bind to ADP-Pi Arp2/3 complexes at branch junctions (Fig. 6). Interestingly, for ADP-Arp2/3 complexes, the protective activity of cortactin provides a means to maintain branched network integrity upon stress application.

In the presence of high ATP concentration, GMF is unable to efficiently prevent branch renucleation, even in the presence of a pulling force (Fig. 5). Actually, GMF could act as a filament producer in cells, by quickly pruning branches generated by the same Arp2/3 complex that remains locally active for several rounds of branch dissociation and regrowth. One can think of GMF as the ‘producer’ of short the actin oligomers that have been reported to be key in lamellipodia turnover (*54*). GMF could also help during cortex regrowth in blebs, by providing the first filaments that invade the bleb, attach to the plasma membrane, and act as seeds to regrow a cortex locally (*55*).

Mechanical stress induces faster debranching. Debranching is accelerated 10 fold for every increase of force by 2 pN (Fig. 1). Any significant stress would thus lead to a change in the network architecture, by changing the network branch connectivity, but also by generating new filaments through branch renucleation. This mechanically-tuned response would be local and would not rely on the presence of membrane-bound NPFs to initiate new branches. Alternatively, at ‘high’ mechanical stress though, Arp2/3 complex leaves with the daughter filament upon debranching, there is no renucleation. This mechanism could potentially be used to quickly disrupt the architecture of a network under high stress.

Future studies should explore the role of mechanical stress and inorganic phosphate on the competition between cortactin and various branch destabilizers, such as ADF/cofilin (*48*), GMF (*37*) and coronin (*39*). This would allow us to better understand how branched actin networks are regulated in a more complex physiological context. Moreover, Arp2/3 complexes exist in different flavors (*18*), thanks to the various combinations of the isoforms of ArpC1A/ArpC1B, ArpC5/ArpC5L and Arp3/Arp3B (*56*), and to prost-translational modifications. Different branch flavors are differentially created depending on sub-cellular localizations and cell types. The diversity of actin networks created in cells is based on a myriad of mecano-chemical signals that finely regulate the assembly and interconnection of actin filaments. This large array of regulation calls for more studies on the fascinating role of the Arp2/3 complex in shaping actin networks.

## Author contributions

JX performed and analysed all experiments RP purified actin and bovine Arp2/3 complexes. AJ and GRL designed and supervised the project, and wrote the manuscript. All authors proofread the manuscript.

## Material and Methods

### Proteins

Alpha skeletal muscle actin (UniProt P68135) was purified from rabbit muscle acetone powder following the protocol from (*57*) as described in (*58*). Actin was fluorescently labeled on the surface accessible lysines using Alexa Fluor 488, Alexa Fluor 568, or Alexa Fluor 647 NHS ester (Thermo Fisher Scientific) as described in (*58*), reaching a final labelling fraction between 15 to 45%.

The Arp2/3 complex was purified from bovine thymus, following a protocol adapted from (*59*).

Spectrin-actin seeds from human red blood cells were purified as described in (*60*).

Recombinant N-terminal glutathione S-transferase (GST) tag human N-WASP-VCA (amino acids 392 to 505, UniProt O00401) were expressed and purified as described in (*61*).

Recombinant N-terminal 6xHis-tagged human GMFγ (UniProt O60234, 7-140 aa from full sequence, obtained from AddGene plasmid #39037) was expressed and purified as described in (*61*).

Recombinant N-terminal GST-tagged mouse cortactin full length (Uniprot Q60598, 1-546 aa) was purified as described in (*15*).

Recombinant human profilin1 (UniProt P07737) was expressed and purified as described in (*62*).

### Buffers

Microfluidics experiments were performed in standard F-buffer containing 5 mM Tris-HCl at pH 7.0, 1 mM MgCl_2_, 0.2 mM EGTA, 0.2 mM ATP (except for experiments with GMF in figure 5C, where it is 2 mM), 10 mM DTT, 1 mM DABCO, supplemented with 0.1% bovine serum albumin (BSA), and 50 mM KCl. For inorganic phosphate experiments, F-buffer was supplemented with KH_2_PO_4_/K_2_HPO_4_ to reach the desired phosphate concentration at pH 7.0, and the concentration of KCl was adjusted to reach a final ionic concentration of 81 mM. BeFx solutions were prepared by mixing 2 mM BeSO_4_ and 10 mM NaF in an F-buffer containing 50 mM KCl.

‘Open’ chambers experiments were performed using F-buffer with 50 mM KCl, supplemented with 0.5 % methylcellulose 4,000 cP.

### Data acquisition

Experiments were performed using a Nikon TiE inverted microscope equipped with a 60× oil-immersion objective, a Kinetix22 SCMOS camera (Photometrics), and a Total Internal Reflection Fluorescence (TIRF) illumination setup (iLAS2, Gataca systems) with 100 mW 488-, 561-, and 642-nm tunable lasers. The temperature was maintained in chamber assay at 25 (± 0.2) °C using a collar objective heater (Okolab). The setup was controlled using micromanager (*63*).

### Microfluidics experiments

Microfluidics experiments were conducted in polydimethylsiloxane (PDMS; Sylgard) chambers based on the original protocol from (*20*), described in detail in (*26*). First, spectrin-actin seeds were attached to the glass surface by flowing in a solution containing 5 pM in F-buffer, for 4 minutes. Next, the surface was passivated by exposing it to a solution containing 5% BSA for 20 minutes. Surface-anchored mother filaments were polymerized by flowing in 0.6 μM 10% Alexa Fluor 488–labeled G-actin for 5 to 10 min. Next, the nucleation of actin filament branches was triggered by flowing in a solution of 20 nM Arp2/3 complex, 50 nM VCA, and 0.4 μM 10% Alexa Fluor 568–labeled G-actin, for around 75 s to obtain a branch density ∼ 1 branch every 10 µm along mother filaments. For experiments in Fig. 1, actin filament branches and mother filaments were aged with a low flow of 0.2 μM 10% Alexa Fluor 568–labeled G-actin for 30 minutes (or 4 minutes for supp Fig. 1). Finally, actin filaments were exposed to debranching conditions. For constant force pulling experiments, we used a solution containing 0.1 μM 10% Alexa Fluor 568–labeled G-actin in different buffer conditions depending, in the presence or absence of GMF or cortactin, on the assay, with an acquisition frame rate of 1 image every 0.5 second to 10 seconds. Please refer to the legend of each corresponding figure for a description of the buffer and protein compositions.

For cortactin experiments, we chose to use 20 nM as our standard concentration for cortactin to be at saturating activity, as McGuirk and colleagues reported an apparent binding affinity of cortactin for Arp2/3 complex at branch junction of 1.3 nM (*37*).

### ‘Open’ chamber experiments

To measure branch detachment without applying any pulling force, we prepared ‘open’ chambers by melting parafilm stripes sandwiched in between cleaned coverslips. This creates chambers of around 10 µL. Chambers are first passivated with 5% BSA solution for 10 minutes, then extensively rinsed with F-buffer with 50 mM KCl. Mother filaments are first assembled in a tube for 10 minutes at room temperature, using 5 µM 10% Alexa Fluor 488–labeled G-actin in F-buffer, with 50 mM KCl. Second, 1.25 µM of preformed mother filaments are next injected inside the chamber in a buffer containing 100 nM Arp2/3, 200 nM VCA, 0.4 µM 10% Alexa Fluor 568–labeled G-actin, 0.4 µM profilin, and supplemented with 0.25 % Methylcellulose. We let the branching reaction proceed for one minute. The branching solution is then replaced by flowing in a solution containing 0.04 µM 10% Alexa Fluor 568–labeled G-actin F-buffer, 50 mM KCl and 0.5 % Methylcellulose. Images are acquired at a frame rate of 1 frame every 30 seconds in TIRF for a total duration of 30 minutes.

## Data analysis

### Fraction of surviving Branches

The fraction of surviving branches as a function of time is computed by the Kaplan-Meier method using the ‘lifelines’ package from python. The error bars represent the standard error calculated by the method of Greenwood.

### Branch renucleation ratio

The branch renucleation ratio represents the ratio of the number of renucleated branches to the number of dissociated branches. Error bars show the binomial Standard deviation.

### Force applied to the branch junction

The tensile force exerted on branches is a result of the viscous drag applied by the fluid to the filaments, as characterized previously in (*21*) for actin filaments. The pulling force on the branch junction is the product of the length of the branch, the local flow velocity, and the longitudinal friction coefficient per unit length of the actin filament (η_actin_ = 6 × 10^−4^ pN.μm^−2^.s). In each experiment, the average force and standard deviation of the force were determined for the population of analyzed branches. In experiments where the force varies over time, we recorded the force at the debranching time for each individual branch to compute the average force of the distribution.

### Affinity constant between Inorganic Phosphate and ADP-Arp2/3 complex at the branch junction

The affinity constant of Pi for the ADP-Arp2/3 complex at a branch junction was derived fitting the observed debranching rates measured at 3.3 pN as a function of phosphate concentration, using the following equation k_deb._ ([Pi]) = k_max_ + (k_min_-k_max_).([Pi]/(K_D_+[Pi])), with k_min_, k_max_ and K_D_ as free parameters. The numerical fit was performed using the ‘curve_fit’ function from the Scipy python package.

### Affinity constant between Inorganic Phosphate and surviving ADP-Arp2/3 complex on mother filament

The affinity constant of Pi for the surviving ADP-Arp2/3 complex that remains attached to the mother filament was estimated by considering the binding follows a simple saturation behaviour, renuc([Pi]) = renuc_ADP_ + (renuc_ADP-BeFx_-renuc_ADP_).([Pi]/(K_D_+[Pi])), with K_D_ as the only free parameters, and considering that the renucleation ratio for ADP-BeFx-Arp2/3 complex branch junctions corresponds to the renucleation ratio that would be obtained at saturating phosphate concentration. From the renucleation ratio obtained at 50 mM Pi, which is half way through the one observed for ADP-Arp2/3 and ADP-BeFx-Arp2/3, we derived that K_D_ to be close to 50 mM.

Note that, according to our reaction scheme, once Pi has departed, the surviving Arp2/3 complex is in the ADP state, from which the reloading of ATP in the presence 200 µM (our standard condition) is much faster (> 1000 s^-1^) and would thus prevent Pi to bind back into the surviving ADP-Arp2/3 complex. As a consequence Pi is not in rapid equilibrium with the surviving ADP-Arp2/3 complex anymore.

### Departure rate of surviving ADP-BeFx-Arp2/3 complex from mother filament

The off-rate of the surviving ADP-BeFx-Arp2/3 complex (Fig. 4C) was derived from the exponential fit of the renucleation ratio of ADP-BeFx-Arp2/3 complexes on mother filaments over time, with an initial value a t = 0 second set at 0.88, which is the average renucleation ratio value obtained in the presence of 1.5 µM actin, as shown in Fig. 4B. Fit was done using the ‘curve_fit function’ from the Scipy python package.

## Supplementary Figures

**figure S1.**
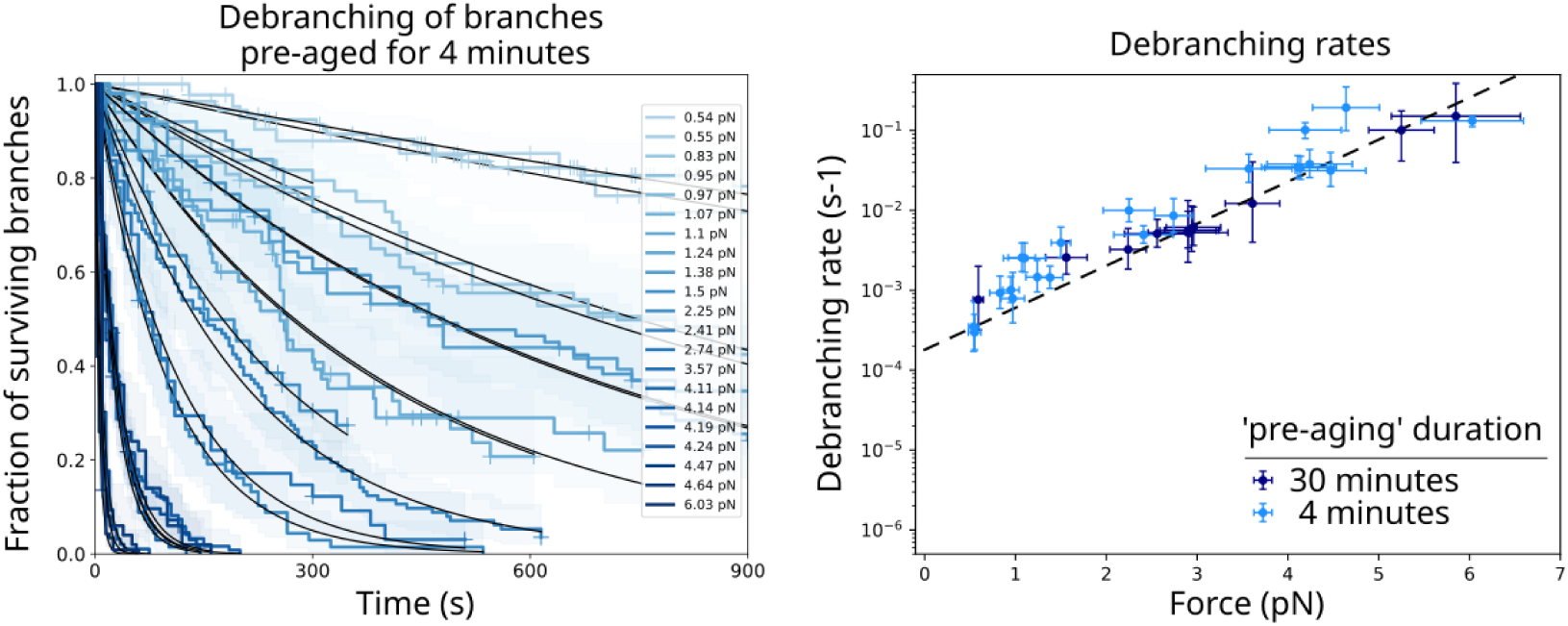
Comparison between 4 and 30 minutes aging. (left) Surviving fractions of branches monitored over time using different flow rates to apply different constant forces, while flowing in 0.15 µM Alexa Fluor 568 (10%)–G-ATP-actin. Black curves are single exponential fits. The average force for each experiment is indicated. Each curve is from a single experiment with at least 50 analyzed branches. Shaded areas are 95% confidence intervals. Vertical ticks on the surviving curves are times where censoring events occurred. (right) Debranching rates as a function of constant pulling forces, for ADP-Arp2/3 complex branch junctions, pre-aged for 4 (light blue) or 30 (blue) minutes. Error bars are standard deviations for the force, and confidence intervals on the single exponential fits, obtained by fitting the upper and lower bounds of 95% confidence intervals of the survival fractions of debranching experiments. Data obtained from experiments with at least 50 and 18 analyzed branches for 4 and 30 minutes pre-aging duration, respectively. Dashed line is a single exponential fit of the 30 minutes pre-aging data.

**figure S2.**
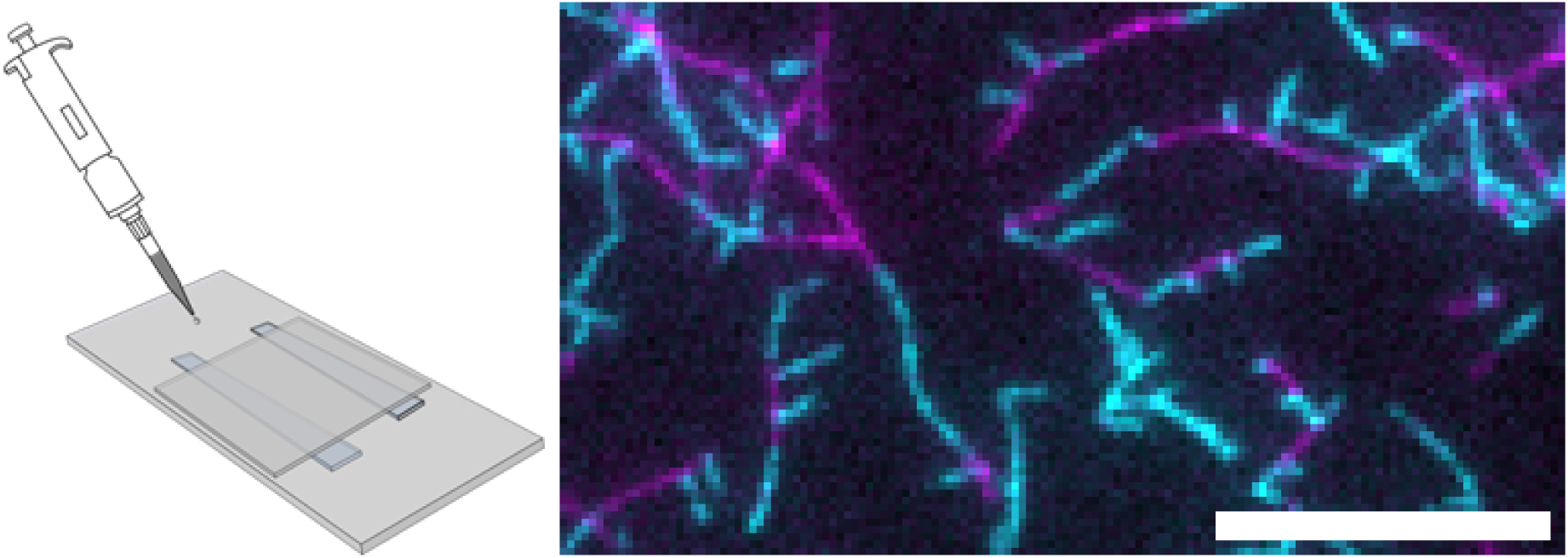
‘Open’ chamber experiment to measure debranching in the absence of force. Branches (cyan) on mother filaments (magenta) are exposed to a solution of 0.04 µM 10% Alexa Fluor 568–labeled G-actin F-buffer, 50 mM KCl and 0.5 % Methylcellulose. Images are acquired at a frame rate of 1 frame every 30 seconds in TIRF for 30 minutes. The scale bar is 10 µm.

**figure S3.**
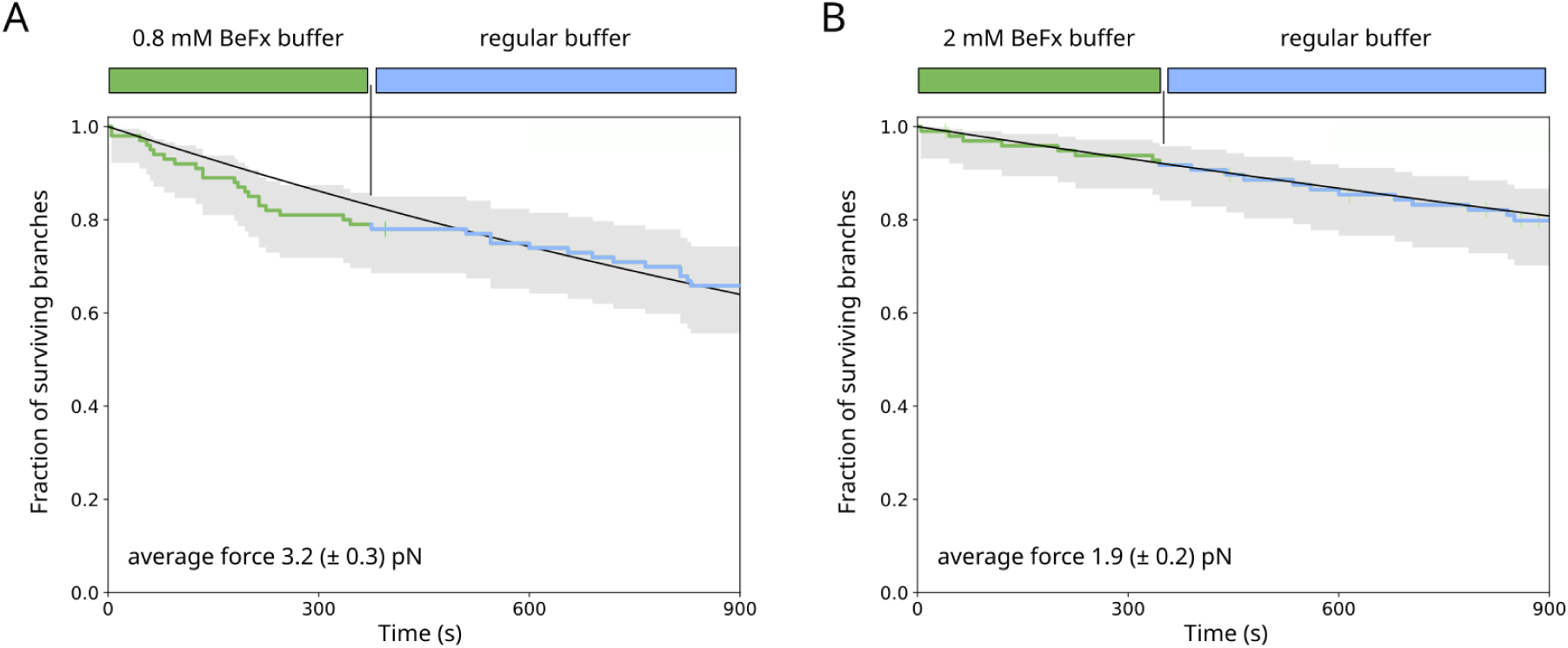
BeFx exposure does not affect debranching rate. Branches initiated as detailed in Fig. 1A are exposed to a solution containing either 0.8 (A) or 2 (B) mM BeFx for 4 minutes. Debranching events are recorded first in the same 0.8 or 2 mM BeFx buffer, before switching to a regular buffer solution. All buffer conditions contain 0.1 µM Alexa Fluor 568 (10%)–G-ATP-actin. For each condition, 100 branches were analyzed. Image acquisition rate is 1 frame every 5 seconds.

**figure S4.**
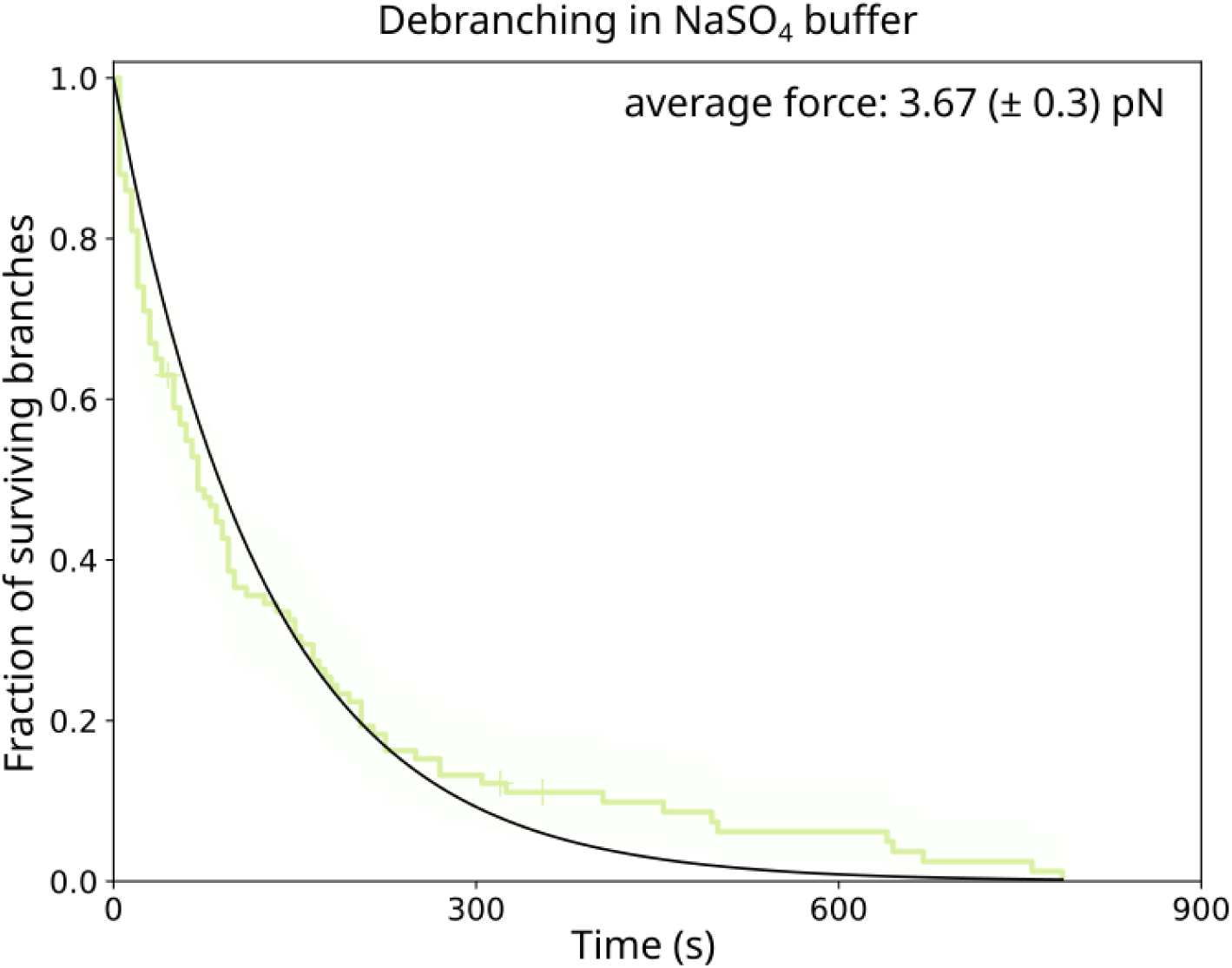
Impact of NaSO_4_ on debranching. In a microfluidics assay, ADP-Arp2/3 complex branch junctions are exposed to a F-buffer supplemented with 0.8 mM NaSO_4_, 23.4 mM NaF and 50 mM KCl, with an average pulling force of 3.67 (± 0.3) pN. The debranching rate is 7.9 10^-3^ (± 0.3) s^-1^. The acquisition time interval is 5 seconds. Number of analyzed branches = 100.

**figure S5.**
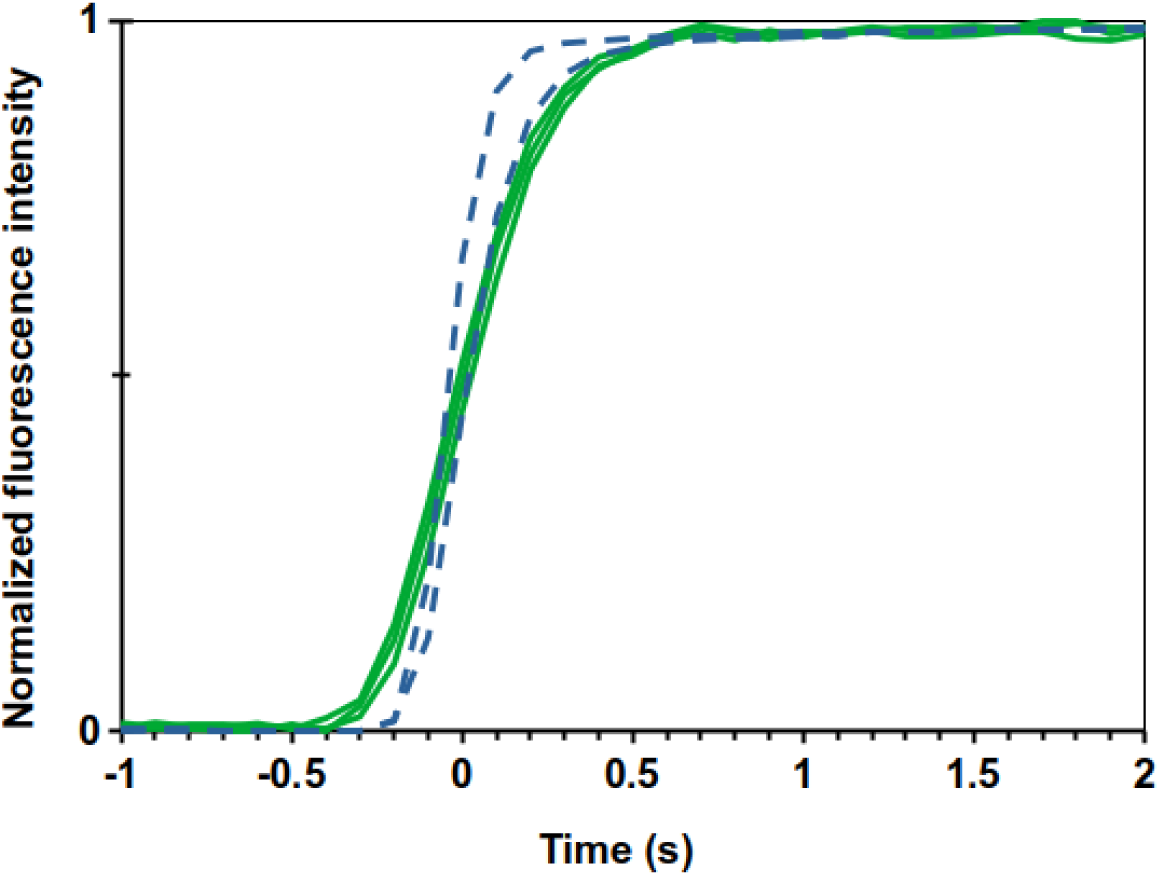
Fast switching between two buffer conditions in microfluidics. Increase in fluorescence intensity upon flow switching in a microfluidics assay, with a flow rate of 10 000 nL/min. Depending on the size of the fluorescent compounds, either Alexa Dye (blue, dashed line) or Alexa-labelled actin (green), the switching is typically between 0.5 and 1 s. Fluorescence images were recorded at 10 images per second in TIRF mode, with an evanescent penetration depth of 150 nm. All curves are positioned with time 0 when the normalized intensity reaches 0.5.

**figure S6.**
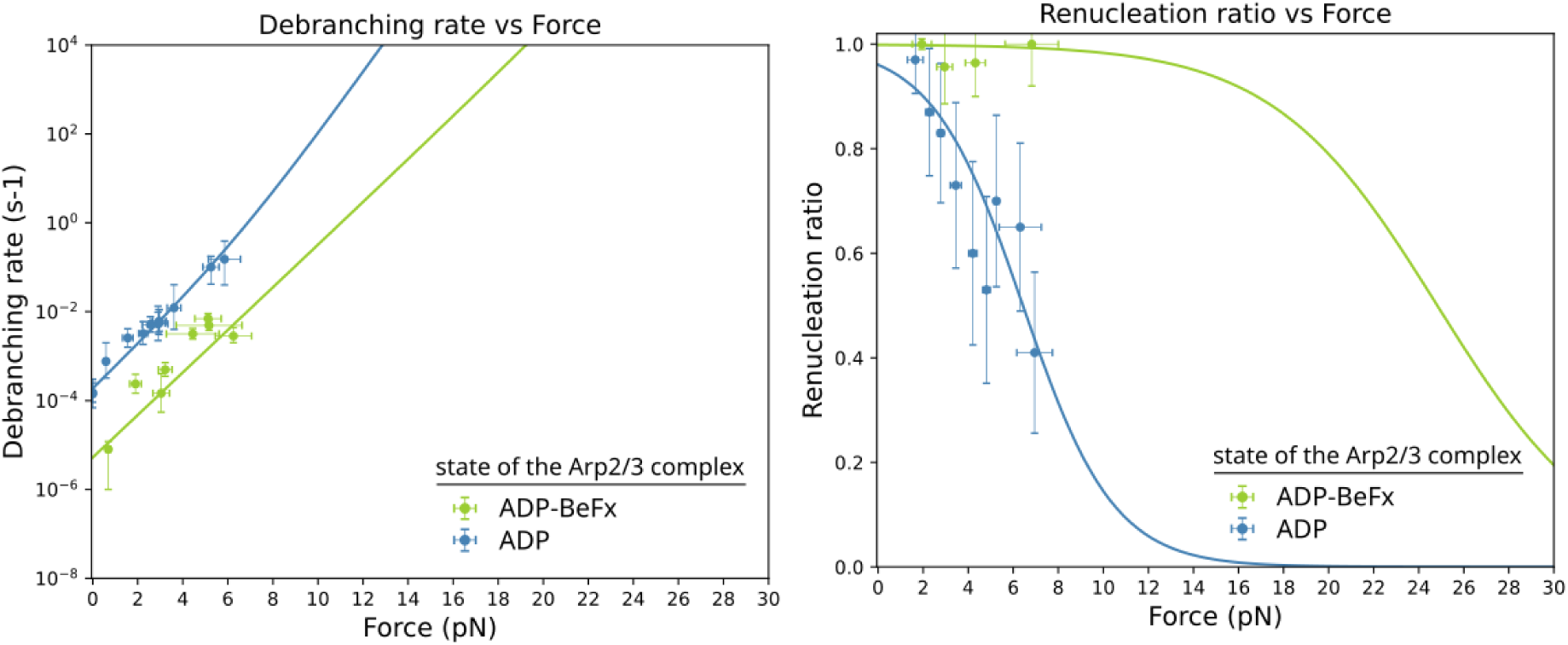
Simultaneous fit of debranching rate and branch renucleation ratio. Same data and fit as in figure 3C,D, displayed from 0 to 30 pN.

**figure S7.**
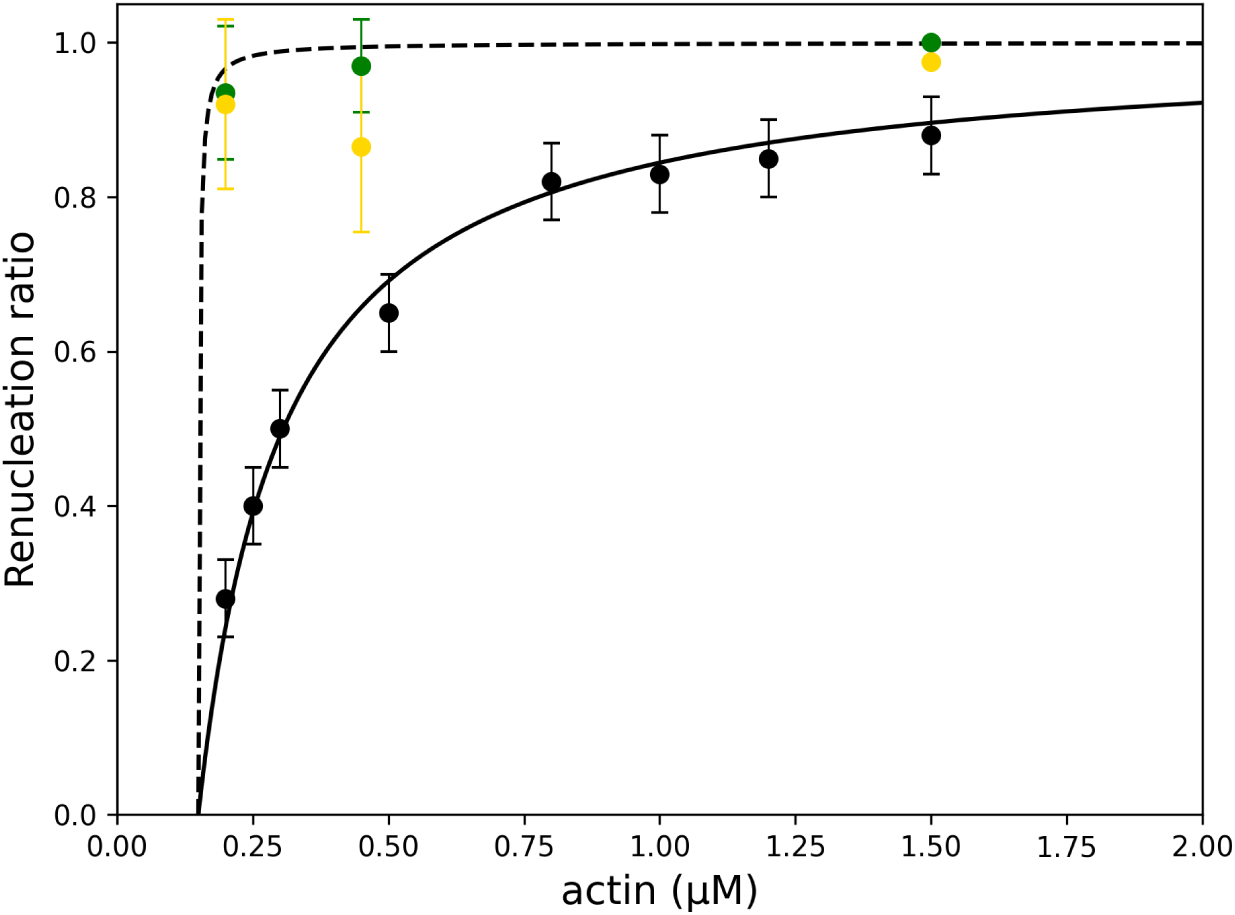
Branch renucleation at different actin concentrations. Branch renucleation ratio from ADP-BeFx-Arp2/3 branches (green) in the presence of 0.2 (n=31 branches), 0.45 (n=32 branches), 1.5 µM actin (n=32 branches), 200 µM ATP, with an average pulling force of 2 pN. Branch renucleation ratio from ADP-Arp2/3 complex branch junctions exposed to 20 nM cortactin (gold), in the presence of 0.2 (n=25 branches), 0.45 (n=31 branches), 1.5 µM actin (n=40 branches), 200 µM ATP, with an average pulling force of 2.2 pN. Black line and data points are from (Ghasemi et al, 2024), and were obtained for 1 pN pulling force. The dashed line corresponds to the theoretical renucleation ratio from ADP-BeFx-Arp2/3 complex, with an actin binding rate to the barbed end of Arp2/3 assumed to be k_on_= 3.4 µM^-1^.s^-1^, in competition with the spontaneous dissociation of surviving ADP-BeFx-Arp2/3 complexes from mother filaments occurring at a rate of 1 10^-4^ s^-1^ (Fig. 4).

**figure S8.**
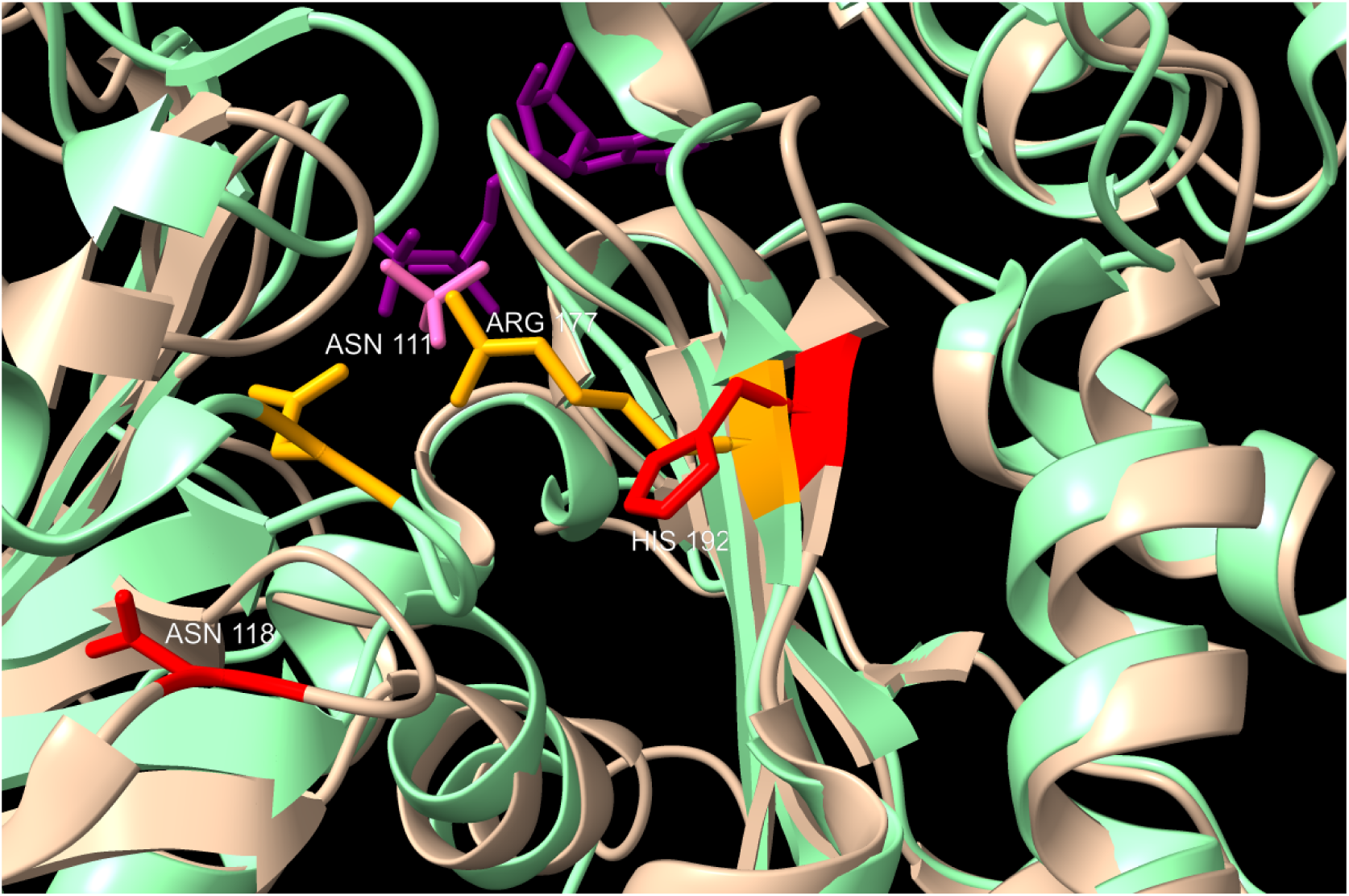
possible Pi backdoor in Arp3 is wide open. Superimposed structures of Arp3 (brown, from PDB 8TAH, (*4*)) and actin (green, from PDB 8A2S, (*25*)), with ADP (purple from both structures) and PO_4_^3-^ (light purple, from actin). Highlighted residues: N111 and R177 from actin in orange, N118 and H192 from Arp3 in red.

## Bibliography

1. A. M. Gautreau, F. E. Fregoso, G. Simanov, R. Dominguez, Nucleation, stabilization, and disassembly of branched actin networks. Trends Cell Biol. 32, 421–432 (2022).

2. T. E. B. Stradal, M. Boiero Sanders, P. Bieling, Arp2/3-complex regulation - Novel insights and open questions. Curr. Opin. Cell Biol. 95, 102565 (2025).

3. A. J. Saks, K. R. Barrie, G. Rebowski, R. Dominguez, NPF binding to Arp2 is allosterically linked to the release of ArpC5’s N-terminal tail and conformational changes in Arp2/3 complex. Proc. Natl. Acad. Sci. U. S. A. 122, e2421557122 (2025).

4. T. van Eeuwen, M. Boczkowska, G. Rebowski, P. J. Carman, F. E. Fregoso, R. Dominguez, Transition State of Arp2/3 Complex Activation by Actin-Bound Dimeric Nucleation-Promoting Factor. Proc. Natl. Acad. Sci. U. S. A. 120, e2306165120 (2023).

5. M. Shaaban, S. Chowdhury, B. J. Nolen, Cryo-EM reveals the transition of Arp2/3 complex from inactive to nucleation-competent state. Nat. Struct. Mol. Biol. 27, 1009–1016 (2020).

6. T. Oda, M. Iwasa, T. Aihara, Y. Maéda, A. Narita, The nature of the globular-to fibrous-actin transition. Nature 457, 441–445 (2009).

7. M. J. Dayel, E. A. Holleran, R. D. Mullins, Arp2/3 complex requires hydrolyzable ATP for nucleation of new actin filaments. Proc. Natl. Acad. Sci. U. S. A. 98, 14871–14876 (2001).

8. C. Le Clainche, D. Pantaloni, M.-F. Carlier, ATP hydrolysis on actin-related protein 2/3 complex causes debranching of dendritic actin arrays. Proc. Natl. Acad. Sci. U. S. A. 100, 6337–6342 (2003).

9. M. J. Dayel, R. D. Mullins, Activation of Arp2/3 complex: addition of the first subunit of the new filament by a WASP protein triggers rapid ATP hydrolysis on Arp2. PLoS Biol. 2, E91 (2004).

10. A. C. Martin, M. D. Welch, D. G. Drubin, Arp2/3 ATP hydrolysis-catalysed branch dissociation is critical for endocytic force generation. Nat. Cell Biol. 8, 826–833 (2006).

11. E. Ingerman, J. Y. Hsiao, R. D. Mullins, Arp2/3 complex ATP hydrolysis promotes lamellipodial actin network disassembly but is dispensable for assembly. J. Cell Biol. 200, 619–633 (2013).

12. B. Ding, H. Y. Narvaez-Ortiz, Y. Singh, G. M. Hocky, S. Chowdhury, B. J. Nolen, Structure of Arp2/3 complex at a branched actin filament junction resolved by single-particle cryo-electron microscopy. Proc. Natl. Acad. Sci. U. S. A. 119, e2202723119 (2022).

13. S. Z. Chou, M. Chatterjee, T. D. Pollard, Mechanism of actin filament branch formation by Arp2/3 complex revealed by a high-resolution cryo-EM structure of the branch junction. Proc. Natl. Acad. Sci. U. S. A. 119, e2206722119 (2022).

14. F. Fäßler, G. Dimchev, V.-V. Hodirnau, W. Wan, F. K. M. Schur, Cryo-electron tomography structure of Arp2/3 complex in cells reveals new insights into the branch junction. Nat. Commun. 11, 6437 (2020).

15. T. Liu, L. Cao, M. Mladenov, A. Jegou, M. Way, C. A. Moores, Cortactin stabilizes actin branches by bridging activated Arp2/3 to its nucleated actin filament. Nat. Struct. Mol. Biol., doi: 10.1038/s41594-023-01205-2 (2024).

16. N. G. Pandit, W. Cao, J. Bibeau, E. M. Johnson-Chavarria, E. W. Taylor, T. D. Pollard, E. M. De La Cruz, Force and phosphate release from Arp2/3 complex promote dissociation of actin filament branches. Proc. Natl. Acad. Sci. U. S. A. 117, 13519–13528 (2020).

17. P. Lappalainen, T. Kotila, A. Jégou, G. Romet-Lemonne, Biochemical and mechanical regulation of actin dynamics. Nat. Rev. Mol. Cell Biol., doi: 10.1038/s41580-022-00508-4 (2022).

18. L. Cao, M. Way, The stabilization of Arp2/3 complex generated actin filaments. Biochem. Soc. Trans. 52, 343–352 (2024).

19. F. Ghasemi, L. Cao, M. Mladenov, B. Guichard, M. Way, A. Jégou, G. Romet-Lemonne, Regeneration of actin filament branches from the same Arp2/3 complex. Sci Adv 10, eadj7681 (2024).

20. A. Jégou, T. Niedermayer, J. Orbán, D. Didry, R. Lipowsky, M.-F. Carlier, G. Romet-Lemonne, Individual actin filaments in a microfluidic flow reveal the mechanism of ATP hydrolysis and give insight into the properties of profilin. PLoS Biol. 9, e1001161 (2011).

21. A. Jégou, M.-F. Carlier, G. Romet-Lemonne, Formin mDia1 senses and generates mechanical forces on actin filaments. Nat. Commun. 4, 1883 (2013).

22. C. Combeau, M. F. Carlier, Probing the mechanism of ATP hydrolysis on F-actin using vanadate and the structural analogs of phosphate BeF-3 and A1F-4. J. Biol. Chem. 263, 17429–17436 (1988).

23. J. Orbán, D. Lorinczy, G. Hild, M. Nyitrai, Noncooperative stabilization effect of phalloidin on ADP.BeFx- and ADP.AlF4-actin filaments. Biochemistry 47, 4530–4534 (2008).

24. A. Muhlrad, P. Cheung, B. C. Phan, C. Miller, E. Reisler, Dynamic properties of actin. Structural changes induced by beryllium fluoride. J. Biol. Chem. 269, 11852–11858 (1994).

25. W. Oosterheert, B. U. Klink, A. Belyy, S. Pospich, S. Raunser, Structural basis of actin filament assembly and aging. Nature, doi: 10.1038/s41586-022-05241-8 (2022).

26. H. Wioland, F. Ghasemi, J. Chikireddy, G. Romet-Lemonne, A. Jégou, Using microfluidics and fluorescence microscopy to study the assembly dynamics of single actin filaments and bundles. J. Vis. Exp., doi: 10.3791/63891 (2022).

27. I. Fujiwara, D. Vavylonis, T. D. Pollard, Polymerization kinetics of ADP- and ADP-Pi-actin determined by fluorescence microscopy. Proc. Natl. Acad. Sci. U. S. A. 104, 8827–8832 (2007).

28. R. Melki, S. Fievez, M.-F. Carlier, Continuous Monitoring of PiRelease Following Nucleotide Hydrolysis in Actin or Tubulin Assembly Using 2-Amino-6-mercapto-7-methylpurine Ribonucleoside and Purine-Nucleoside Phosphorylase as an Enzyme-Linked Assay†. [Preprint] (1996). 10.1021/bi961325o.

29. W. Oosterheert, F. E. C. Blanc, A. Roy, A. Belyy, M. B. Sanders, O. Hofnagel, G. Hummer, P. Bieling, S. Raunser, Molecular mechanisms of inorganic-phosphate release from the core and barbed end of actin filaments. Nat. Struct. Mol. Biol., doi: 10.1038/s41594-023-01101-9 (2023).

30. A. Schahl, L. Lagardère, B. Walker, P. Ren, H. Wioland, M. Ballet, A. Jégou, M. Chavent, J.-P. Piquemal, Histidine 73 methylation coordinates β-actin plasticity in response to key environmental factors. Nat. Commun. 16, 1–13 (2025).

31. O. K. Dudko, G. Hummer, A. Szabo, Theory, analysis, and interpretation of single-molecule force spectroscopy experiments. Proc. Natl. Acad. Sci. U. S. A. 105, 15755–15760 (2008).

32. B. L. Goode, M. O. Sweeney, J. A. Eskin, GMF as an Actin Network Remodeling Factor. Trends Cell Biol., doi: 10.1016/j.tcb.2018.04.008 (2018).

33. M. Poukkula, M. Hakala, N. Pentinmikko, M. O. Sweeney, S. Jansen, J. Mattila, V. Hietakangas, B. L. Goode, P. Lappalainen, GMF promotes leading-edge dynamics and collective cell migration in vivo. Curr. Biol. 24, 2533–2540 (2014).

34. Q. Luan, B. J. Nolen, Structural basis for regulation of Arp2/3 complex by GMF. Nat. Struct. Mol. Biol. 20, 1062–1068 (2013).

35. M. Boczkowska, G. Rebowski, R. Dominguez, Glia Maturation Factor (GMF) Interacts with Arp2/3 Complex in a Nucleotide State-dependent Manner. J. Biol. Chem. 288, 25683–25688 (2013).

36. L. Blanchoin, T. D. Pollard, Mechanism of interaction of Acanthamoeba actophorin (ADF/Cofilin) with actin filaments. J. Biol. Chem. 274, 15538–15546 (1999).

37. E. R. McGuirk, N. Koundinya, P. Nagarajan, S. B. Padrick, B. L. Goode, Direct observation of cortactin protecting Arp2/3-actin filament branch junctions from GMF-mediated destabilization. Eur. J. Cell Biol. 103, 151378 (2024).

38. C. A. Ydenberg, S. B. Padrick, M. O. Sweeney, M. Gandhi, O. Sokolova, B. L. Goode, GMF severs actin-Arp2/3 complex branch junctions by a cofilin-like mechanism. Curr. Biol. 23, 1037–1045 (2013).

39. N. Koundinya, R. M. Aguilar, K. Wetzel, M. R. Tomasso, P. Nagarajan, E. R. McGuirk, S. B. Padrick, B. L. Goode, Two ligands of Arp2/3 complex, yeast coronin and GMF, interact and synergize in pruning branched actin networks. J. Biol. Chem. 0, 108191 (2025).

40. M. Schnoor, T. E. Stradal, K. Rottner, Cortactin: Cell Functions of A Multifaceted Actin-Binding Protein. Trends Cell Biol. 28, 79–98 (2018).

41. L. A. Helgeson, B. J. Nolen, Mechanism of synergistic activation of Arp2/3 complex by cortactin and N-WASP. Elife 2, e00884 (2013).

42. L. A. Helgeson, J. G. Prendergast, A. R. Wagner, M. Rodnick-Smith, B. J. Nolen, Interactions with actin monomers, actin filaments, and Arp2/3 complex define the roles of WASP family proteins and cortactin in coordinately regulating branched actin networks. J. Biol. Chem. 289, 28856–28869 (2014).

43. F. E. Fregoso, M. Boczkowska, G. Rebowski, P. J. Carman, T. van Eeuwen, R. Dominguez, Mechanism of synergistic activation of Arp2/3 complex by cortactin and WASP-family proteins. Nat. Commun. 14, 6894 (2023).

44. B. A. Smith, S. B. Padrick, L. K. Doolittle, K. Daugherty-Clarke, I. R. Corrêa Jr, M.-Q. Xu, B. L. Goode, M. K. Rosen, J. Gelles, Three-color single molecule imaging shows WASP detachment from Arp2/3 complex triggers actin filament branch formation. Elife 2, e01008 (2013).

45. S. S. Chavali, S. Z. Chou, W. Cao, T. D. Pollard, E. M. De La Cruz, C. V. Sindelar, Cryo-EM structures reveal how phosphate release from Arp3 weakens actin filament branches formed by Arp2/3 complex. Nat. Commun. 15, 2059 (2024).

46. S. Z. Chou, T. D. Pollard, Mechanism of actin polymerization revealed by cryo-EM structures of actin filaments with three different bound nucleotides. Proc. Natl. Acad. Sci. U. S. A., doi: 10.1073/pnas.1807028115 (2019).

47. T. Paramanathan, D. Reeves, L. J. Friedman, J. Kondev, J. Gelles, A general mechanism for competitor-induced dissociation of molecular complexes. Nat. Commun. 5, 5207 (2014).

48. J. Chung, B. L. Goode, J. Gelles, Single-molecule analysis of actin filament debranching by cofilin and GMF. Proceedings of the National Academy of Sciences 119, e2115129119 (2022).

49. R. J. Gillies, T. Ogino, R. G. Shulman, D. C. Ward, 31P nuclear magnetic resonance evidence for the regulation of intracellular pH by Ehrlich ascites tumor cells. J. Cell Biol. 95, 24–28 (1982).

50. S. Soboll, R. Scholz, H. W. Heldt, Subcellular metabolite concentrations. Dependence of mitochondrial and cytosolic ATP systems on the metabolic state of perfused rat liver. Eur. J. Biochem. 87, 377–390 (1978).

51. K. van Eunen, J. Bouwman, P. Daran-Lapujade, J. Postmus, A. B. Canelas, F. I. C. Mensonides, R. Orij, I. Tuzun, J. van den Brink, G. J. Smits, W. M. van Gulik, S. Brul, J. J. Heijnen, J. H. de Winde, M. J. T. de Mattos, C. Kettner, J. Nielsen, H. V. Westerhoff, B. M. Bakker, Measuring enzyme activities under standardized in vivo-like conditions for systems biology: Standardized enzyme assays for systems biology. FEBS J. 277, 749–760 (2010).

52. M. M. Lacy, D. Baddeley, J. Berro, Single-molecule turnover dynamics of actin and membrane coat proteins in clathrin-mediated endocytosis. Elife 8 (2019).

53. S. I. Mousavi, M. M. Lacy, X. Li, J. Berro, Fast actin disassembly and fimbrin mechanosensitivity support rapid turnover in a model of clathrin-mediated endocytosis. Cytoskeleton (Hoboken), doi: 10.1002/cm.22002 (2025).

54. D. Raz-Ben Aroush, N. Ofer, E. Abu-Shah, J. Allard, O. Krichevsky, A. Mogilner, K. Keren, Actin Turnover in Lamellipodial Fragments. Curr. Biol. 27, 2963–2973.e14 (2017).

55. J. M. García-Arcos, J. Ziegler, S. Grigolon, L. Reymond, G. Shajepal, C. J. Cattin, A. Lomakin, D. J. Müller, V. Ruprecht, S. Wieser, R. Voituriez, M. Piel, Rigidity percolation and active advection synergize in the actomyosin cortex to drive amoeboid cell motility. Dev. Cell, doi: 10.1016/j.devcel.2024.06.023 (2024).

56. J. V. G. Abella, C. Galloni, J. Pernier, D. J. Barry, S. Kjær, M.-F. Carlier, M. Way, Isoform diversity in the Arp2/3 complex determines actin filament dynamics. Nat. Cell Biol. 18, 76–86 (2016).

57. J. A. Spudich, S. Watt, The regulation of rabbit skeletal muscle contraction. I. Biochemical studies of the interaction of the tropomyosin-troponin complex with actin and the proteolytic fragments of myosin. J. Biol. Chem. 246, 4866–4871 (1971).

58. G. Romet-Lemonne, B. Guichard, A. Jégou, Using Microfluidics Single Filament Assay to Study Formin Control of Actin Assembly. Methods Mol. Biol. 1805, 75–92 (2018).

59. H. N. Higgs, L. Blanchoin, T. D. Pollard, Influence of the C terminus of Wiskott-Aldrich syndrome protein (WASp) and the Arp2/3 complex on actin polymerization. Biochemistry 38, 15212–15222 (1999).

60. H. Wioland, B. Guichard, Y. Senju, S. Myram, P. Lappalainen, A. Jégou, G. Romet-Lemonne, ADF/Cofilin Accelerates Actin Dynamics by Severing Filaments and Promoting Their Depolymerization at Both Ends. Curr. Biol. 27, 1956–1967.e7 (2017).

61. L. Cao, F. Ghasemi, M. Way, A. Jégou, G. Romet-Lemonne, Regulation of branched versus linear Arp2/3-generated actin filaments. EMBO J. 42, e113008 (2023).

62. L. Cao, M. Kerleau, E. L. Suzuki, H. Wioland, S. Jouet, B. Guichard, M. Lenz, G. Romet-Lemonne, A. Jegou, Modulation of formin processivity by profilin and mechanical tension. Elife 7, e34176 (2018).

63. A. D. Edelstein, M. A. Tsuchida, N. Amodaj, H. Pinkard, R. D. Vale, N. Stuurman, Advanced methods of microscope control using μManager software. J Biol Methods 1 (2014).

